# Gap junctional coupling of molecular layer interneurons enables transient NMDA driven synchronization

**DOI:** 10.64898/2026.05.28.728326

**Authors:** Nils A. Koch, Anmar Khadra

## Abstract

Molecular layer interneurons (MLIs) play a crucial role in modulating the output of the cerebellar cortex through their inhibition of Purkinje cells. MLIs also inhibit other MLIs synaptically and are coupled electrically through gap junctions. While synchronization of MLIs has been observed, comprehensive understanding of the role of gap junctional coupling in shaping MLI network activity is lacking. Dendro-dendritic gap junctional coupling in MLIs involves propagation of signals to and from the dendritic gap junction location which can lead to neural synchronization. However, how this is regulated by the intrinsic electrical properties of MLIs, including dendritic properties, is poorly understood. In this study, we apply conductance-based computational modelling to examine the effect of dendritic filtering on gap junctional coupling in pairs of ball-and-stick MLI models, demonstrating that gap junctional properties, rather than the active dendritic properties of MLIs, primarily dictate gap junction-driven synchronization. By systematically reducing the ball-and-stick model to a one-compartment MLI model, we additionally investigate the role of MLI gap junctional coupling in mediating MLI network synchrony. Our results reveal that transient AMPA input drives brief network-wide synchronization, whereas NMDA-mediated elevated firing enables gap junction-dependent oscillatory synchronization that is further enhanced by MLI–MLI inhibition in a positive feedback loop, producing pronounced peaks of network coactivity resembling sensory-evoked MLI activity observed *in vivo*. These findings provide important insights into network dynamics of MLIs and how gap junctions shape their activity, with broader implications for other neural networks that rely on gap junctional coupling.

## Introduction

In the cerebellum, molecular layer interneurons (MLIs) have typically been classified into cerebellar stellate cells (CSCs) and basket cells (BCs) based on, respectively, their positions in the distal and proximal thirds of the molecular layer as well as their innervation of Purkinje cell (PC) dendrites and soma (Kim and Augustine, 2021; Brown et al., 2019). However, morphological characterization of MLIs has demonstrated that a continuum of cellular morphologies exists along the proximal-distal axis of the molecular layer (Wang and Lefebvre, 2022), with transcriptomic analysis suggesting that 2 distinct sub-populations of MLIs exist throughout the molecular layer independent of position (Kozareva et al., 2021). These two MLI subgroups, MLI1 and MLI2, have distinct circuit roles: MLI1s inhibit PCs directly, whereas MLI2s inhibit MLI1s thereby disinhibiting PCs (Lackey et al., 2024). Additionally, MLIs collectively have been shown to express the gap junctional protein Connexin 36 (Alcami and Marty, 2013), with such gap junctional connectivity demonstrated through both dye coupling (Mann-Metzer and Yarom, 1999) and the presence of spikelets (Hoehne et al., 2020; Kozareva et al., 2021). Interestingly, a proximal-distal gradient of electrical coupling is observed in the molecular layer, with greater gap junctional connections among BCs compared to CSCs (Alcami and Marty, 2013). However, with respect to the two MLI subgroups, only MLI1s express Connexin 36 and are electrically coupled through dendro-dendritic and dendro-somatic gap junctions (Kozareva et al., 2021; Sotelo and Llinás, 1972), allowing for spikelets to form. However, with respect to the two MLI subgroups, only MLI1s express Connexin 36 and are electrically coupled through dendro-dendritic and dendro-somatic gap junctions (Kozareva et al., 2021; Sotelo and Llinás, 1972), allowing for spikelets to form.

Firing activity that is consistent with gap junctional coupling, such as synchronous firing (Lackey et al., 2024; Blot et al., 2016) and spatially correlated firing (Han et al., 2018), has also been observed in MLIs. Consistent with these observations, MLI electrical coupling contributes to the spread of synaptic events across coupled cells to promote synchrony (Hoehne et al., 2020) and is proposed to contribute to lateral inhibition in the cerebellar cortex through spatial synchronization of MLI inhibition on PCs (Kim and Augustine, 2021). Optogenetic mapping of MLIs suggest that seven MLIs converge onto one PC with gap junctional connectivity in clusters of 3–5 electrically coupled MLIs (Kim et al., 2014; Mann-Metzer and Yarom, 1999; Alcami and Marty, 2013). Pharmacological block of gap junctions reduces this MLI convergence onto PCs (Kim et al., 2014), suggesting that electrical coupling of MLIs enhances PC inhibition. Indeed, *in vivo* voltage imaging revealed large scale transient MLI synchronization in response to sensory input that inhibits Purkinje cells in a manner sufficient to evoke or augment motor responses to the sensory stimulus (Brown et al., 2025). Interestingly, this large transient synchronization of MLIs was followed by elevated activity for hundreds of ms with trials that have high transient synchronization exhibiting damped oscillations in firing rate during this elevated activity (Brown et al., 2025).

Active, or voltage-gated, dendritic conductances contribute to synaptic filtering and firing properties of neurons (Migliore and Shepherd, 2002; Tran-Van-Minh et al., 2015; Gollo et al., 2009). In CSCs, NMDA receptor activation and voltage-gated calcium (Ca^2+^) channels contribute to supralinear dendritic Ca^2+^elevation in response to high frequency input (Tran-Van-Minh et al., 2015). Although sublinear voltage integration has been observed in CSC dendrites and attributed to the passive properties of their thin dendritic processes (Biane et al., 2021; Tran-Van-Minh et al., 2015; Abrahamsson et al., 2012), modelling work in parvalbumin positive GABAergic hippocampal basket cells has shown that active dendritic conductances dictate the extent to which network dynamics are influenced by gap junctional coupling (Saraga et al., 2006); this suggests that active dendritic conductances may also contribute to gap junctional coupling in MLIs. In particular, the requirement of APs to propagate back through dendrites to evoke gap junctional currents across dendro-dendritic gap junctions, followed by the propagation of these signals toward the soma of the coupled cell, may distinguish the role of active dendritic conductances in electrical coupling of MLIs from the sublinear integration of synaptically evoked voltage responses. In agreement with this idea, backpropagating APs activate a number of active dendritic currents in a multi-compartmental MLI conductance-based model (Rizza et al., 2021), as well as in multiple multi-compartmental BC models (Masoli et al., 2025).

In this study, we investigate how dendritic currents affect synchronization in pairs of gap junctionally coupled Hodgkin-Huxley type MLI models and the role of gap junctional connectivity in MLI responses observed *in vivo*, particularly their dendritic filtering properties. Specifically, we demonstrate that gap junctional properties and not active dendritic properties dictate gap junction driven synchronization in pairs of MLI. We also show that, in network models of MLI, NMDA depolarization is required for gap junctionally mediated transient synchronization in these cells.

## Materials and Methods

### Ball and stick MLI model

To investigate spatial aspects involving dendritic filtering, a ball and stick model consisting of a soma (the ball) and a dendritic cable (the stick) was developed using the formulism of Goldberg et al. (2007). The cable equation for the stick with length *L* and radius *r*_*cable*_ is given by

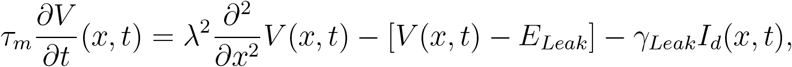

where 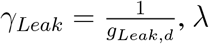 is the membrane length constant, *τ*_*m*_ = *R*_*m*_*C*_*m*_ is the membrane time constant 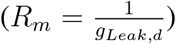, and *I*_*d*_ is the sum of dendritic currents including delayed rectifier K^+^ (*I*_*Kdr*_), A-type K^+^ (*I*_*A*_), Ca^2+^ activate K^+^(*I*_*K*(*Ca*)_), T-type Ca^2+^ (*I*_*T*_ ), high voltage activated (HVA) Ca^2+^ (*I*_*HVA*_) and leak (*I*_*Leak*_) currents, each given by

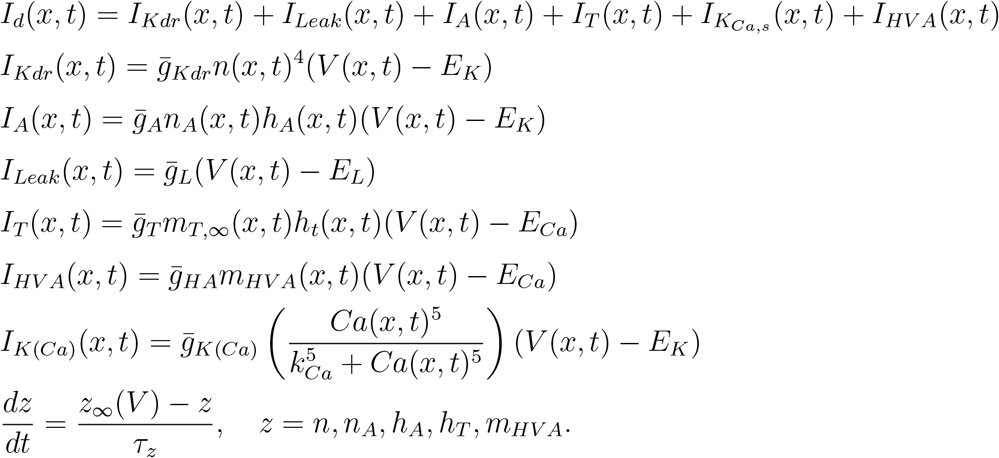

The dendritic stick is discretized using the finite difference method into *n* = 50 dendritic compartments, each with length 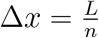, yielding the following equations

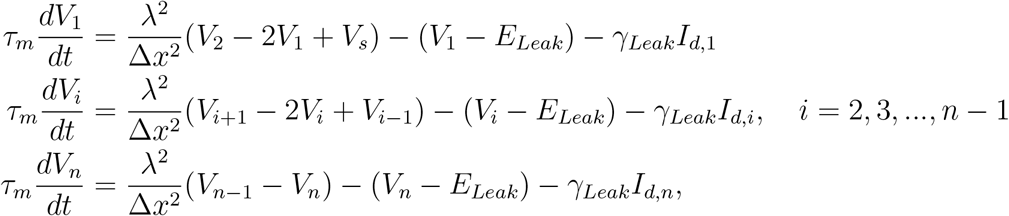

with a sealed end at *x* = *L*.

Ca^2+^ dynamics is governed by the flux-balance equation, given by

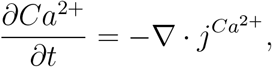

where no-flux boundary conditions are assumed and the flux 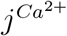 is described by the Nernst-Planck Equation, given by

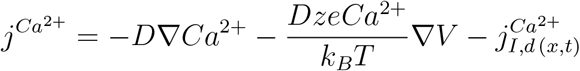

with

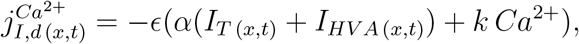

*z* = 2 is the valence of Ca^2+^, *e* is the elementary charge, *k*_*B*_ is the Boltzmann constant, and *T* = 309.15 K is the absolute temperature. The discretized form of the continuity equation is thus given by

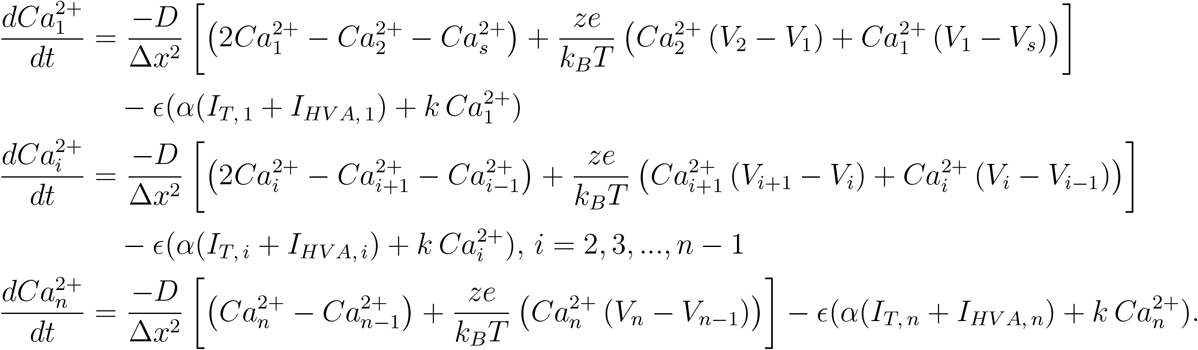

A Hodgkin-Huxley type model is incorporated at *x* = 0 where the ball (of radius *r*_*s*_) representing the soma is located. The dynamic equations representing the soma are modified from Farjami et al. (2020b) and are given by (in a compartmentalized form consistent with the spatial discretization)

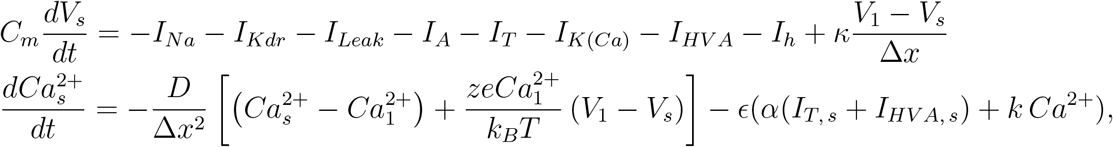

where

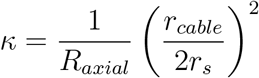

(with *R*_*axial*_ representing the internal resistance of the cable in Ω·cm^2^), *I*_*Na*_ is the somatic Na^+^ current, given by

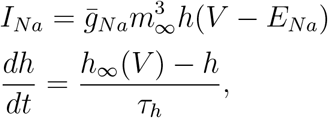

and *I*_*h*_ is the somatic hyperpolarization-activated current, given by (Rizza et al., 2021; Angelo et al., 2007)

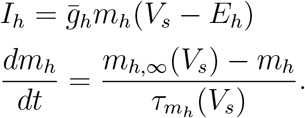

The maximal conductances (Table 1) were fit to *in vitro* recordings (Rizza et al., 2021; Locatelli et al., 2020) with gating parameters modified according to Farjami et al. (2020a) in Table 2 and additional model parameters detailed in Table 3.

**Table 1:**
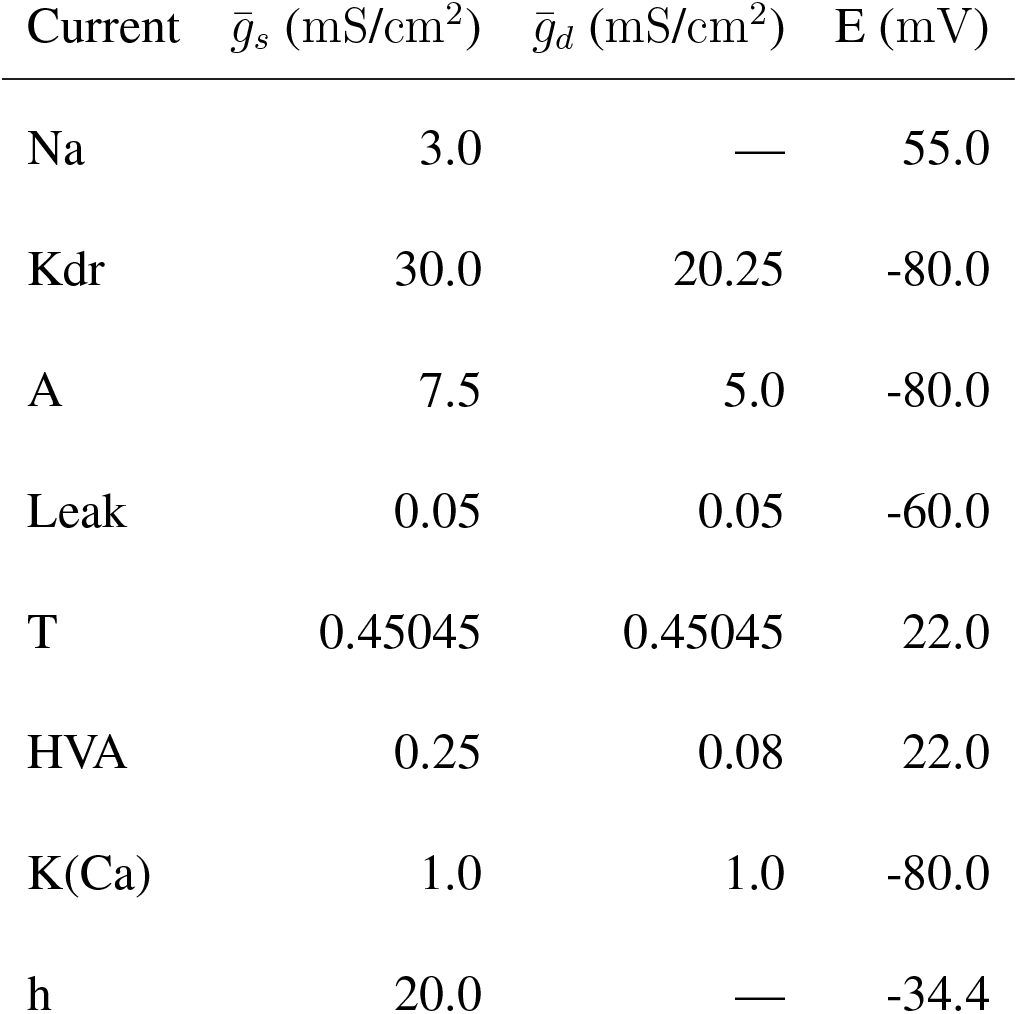
Maximal Conductances for MLI cable model.

**Table 2:**
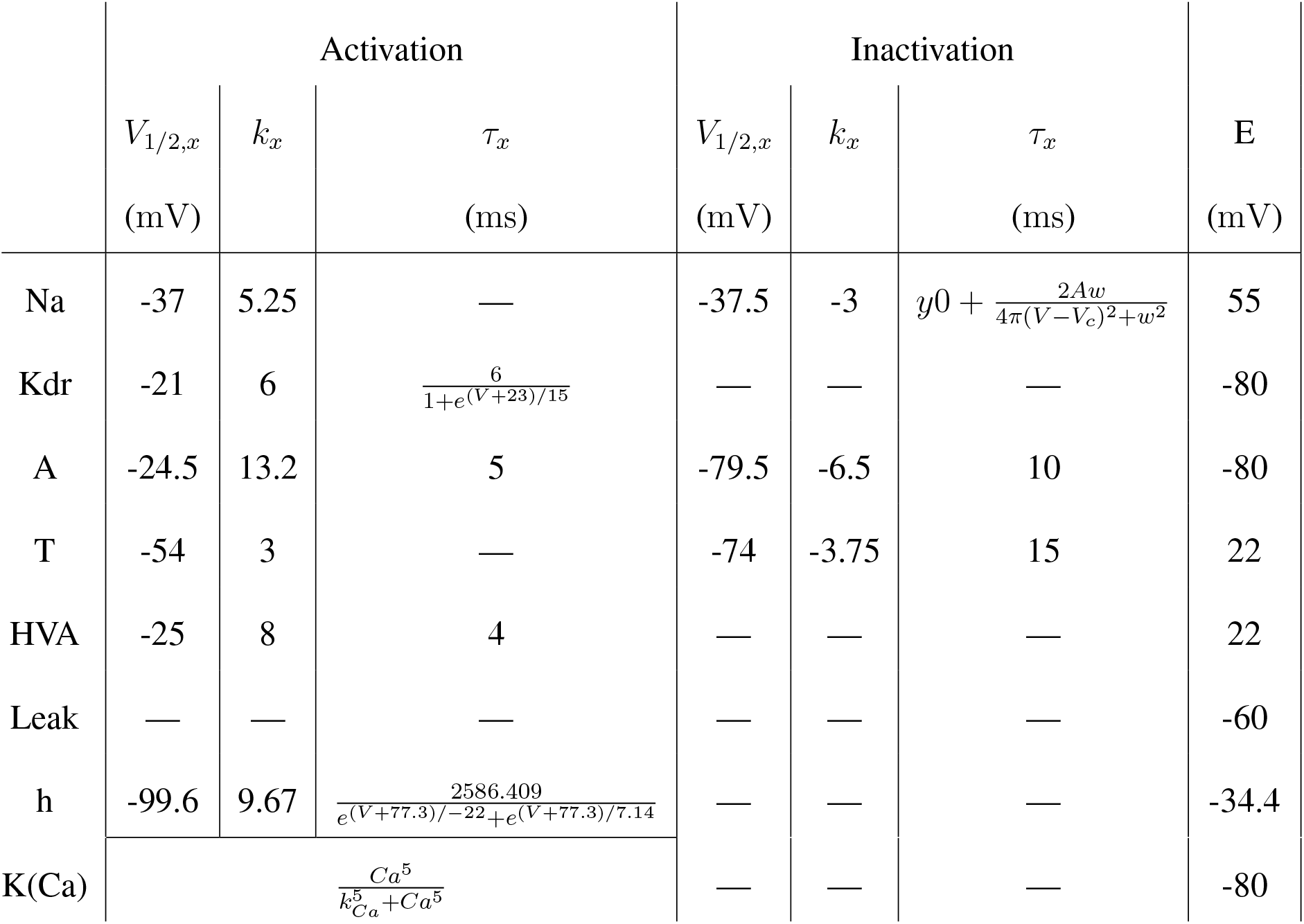
Steady state activation and inactivation are defined as Boltzmann functions of the form 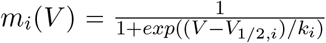 for each current. Time constants for activation and inactivation as well as reversal potentials are listed for each current. Values that are different from those in the one compartment model are listed in parentheses.

|  | Activation |  |  | Inactivation |  |  | E |
| --- | --- | --- | --- | --- | --- | --- | --- |
| | $V_{1/2,x}$<br>(mV) | $k_x$ | $\tau_x$<br>(ms) | $V_{1/2,x}$<br>(mV) | $k_x$ | $\tau_x$<br>(ms) | |
| Na | -37 | 5.25 | — | -37.5 | -3 | $y0 + \frac{2Aw}{4\pi(V-V_c)^2+w^2}$ | 55 |
| Kdr | -21 | 6 | $\frac{6}{1+e^{(V+23)/15}}$ | — | — | — | -80 |
| A | -24.5 | 13.2 | 5 | -79.5 | -6.5 | 10 | -80 |
| T | -54 | 3 | — | -74 | -3.75 | 15 | 22 |
| HVA | -25 | 8 | 4 | — | — | — | 22 |
| Leak | — | — | — | — | — | — | -60 |
| h | -99.6 | 9.67 | $\frac{2586.409}{e^{(V+77.3)/-22}+e^{(V+77.3)/7.14}}$ | — | — | — | -34.4 |
| K(Ca) | $\frac{Ca^5}{k_{Ca}^5+Ca^5}$ | | | — | — | — | -80 |

**Table 3:** Parameter values for MLI cable model.

|  |  |  |
| --- | --- | --- |
| $R_{axial}$ | 110 $\Omega$.cm | Rizza et al. (2021) |
| $r_s$ | 6 $\mu\text{m}$ | Wang and Lefebvre (2022) |
| $r_{cable}$ | 0.2 $\mu\text{m}$ | Abrahamsson et al. (2012) |
| L | 150 $\mu\text{m}$ | Wang and Lefebvre (2022) |
| $C_m$ | 1.50148 $\mu\text{F}/\text{cm}^2$ | Farjami et al. (2020a) |
| $\epsilon$ | 0.015 | Farjami et al. (2020a) |
| $\alpha$ | 0.018 $\mu\text{M cm}^2 \mu\text{A}^{-1} \text{ms}^{-1}$ | Farjami et al. (2020a) |
| w | 46 mV | Farjami et al. (2020a) |
| y0 | 0.1 ms | Farjami et al. (2020a) |
| A | 322 ms.mV | Farjami et al. (2020a) |
| k | 0.1 $\text{ms}^{-1}$ | Farjami et al. (2020a) |
| $k_{Ca}$ | 0.45 $\mu\text{M}$ | Farjami et al. (2020a) |
| D | 5.3 x 10<sup>-9</sup> $\text{cm}^2/\text{ms}$ | — |
| $\lambda$ | 350.33 $\mu\text{m}$ | $\lambda = \sqrt{\frac{r_{cable} R_m}{2R_{axial}}}$ |

### Quantifying dendritic filtering in the ball and stick MLI model

Dendritic filtering properties were assessed in the absence of APs generated through the removal of the somatic sodium current (g_Na_ = 0). To investigate the dendritic filtering occurring to distal synaptic inputs, the distal cable boundary condition is set to an Ornstein-Uhlenbeck (OU) noise process with rate constant *θ* = 0.5 ms^−1^, mean *μ* given by the value of *V*_*n*_ in the distal dendritic compartment when *V*_*s*_ = −50 mV, and noise intensity *σ* = 30. Additionally, a linear chirp stimulus (0.001 – 100 Hz, with amplitudes 5, 10 and 15 mV applied over 1000 s and centred at −50 mV) is applied to the somatic end of the cable to probe the filtering of backpropagating subthreshold somatic membrane potentials under normal conditions. With both input stimuli, dendritic currents are removed individually to assess their impact on dendritic filtering. To quantify filtering, power gain in decibels (dB) is calculated in the frequency domain as the 10log_10_(p), where *p* is the ratio of the power in the recording compartment to the power in the input.

### Gap-junctional coupling in ball and stick MLI models

Gap junctional coupling between MLIs do not display rectification or voltage dependence (Alcami and Marty, 2013). Therefore, linear non-rectifying gap-junctions were incorporated between two cells: *m* and *u*. For a gap junction connecting compartment *i* in cell *m* to compartment *j* in cell *u* (*i, j* = 1, 2, …, *n*), the gap junctional current is given by 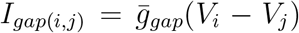. The gap junctional currents *I*_*gap*(*i,j*)_ and *I*_*gap*(*j,i*)_ are added to *I*_*d,i*_ and *I*_*d,j*_ in the discretized cable equations of the ball and stick models of cells *m* and *u*, respectively.

### MLI Network simulations

The synaptic and gap junctional networks of MLIs occur in the sagittal plane and do not extend in the transverse plane (Rieubland et al., 2014; Kim et al., 2014; Palay and Chan-Palay, 1974; Kim and Augustine, 2021). Correspondingly, MLI network models are generated by tiling a 2-dimensional sagittal plane of the molecular layer 3000 μm by 200 μm using the distribution of nearest neighbours in the sagittal plane (Rieubland et al., 2014); the latter is approximated as the probability density function of a log-normal distribution of the form

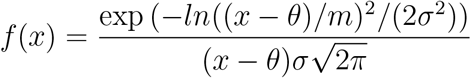

with parameters *θ* = −2.3137, *σ* = 0.5122, and *m* = 49.8240 (Figure S1A) at a density of 28000/mm^3^ (Santos-Valencia et al., 2025) with transverse plane thickness of 20 μm (Rieubland et al., 2014) and a minimum intersomatic distance of 10 μm. This process is repeated to generate 100 different network realizations of similar scale (*M* = 336 interneurons per network) to the maximum simultaneously recorded by Brown et al. (2025).

Connectivity between MLIs in the network is generated probabilistically in a pairwise fashion based on their intersomatic distance and distance dependent probabilities for both chemical and electrical connections (Rieubland et al., 2014). Both of these chemical and electrical distance-dependent probabilities are fit to a gamma probability density function of form

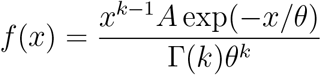

with *k* = 1.41, *θ* = 58.34, and *A* = 26.91 for chemical synapses (Figure S1B cyan), and *k* = 1.81, *θ* = 30.42, and *A* = 44.47 for electrical synapses (Figure S1B, orange). However, chemical synapse probability exhibits a distal-to-proximal gradient, with connections increasingly more prevalent towards proximal positions within the molecular layer (Rieubland et al., 2014). As such, synaptic connections within the network are assigned based on a distance-dependent probability, scaled by a directional probability determined by the location of the interneurons in the molecular layer (Table S1). Likewise, the probability of electrical synapses between MLIs is spatially biased, with MLIs in the third of the molecular layer proximal to the Purkinje cell layer more likely to have gap junctional connections than those in the more distal two thirds of the molecular layer (Rieubland et al., 2014). A distance-dependent probability of gap junctional connectivity is thus also scaled according to the location of the two interneurons in the molecular layer (Table S2).

Because the behaviour of the ball and stick models is governed by the passive dendritic properties, both in terms of dendritic filtering and dominance of gap junctional properties in mediating electrical coupling between pairs of MLIs, such models are replaced with one compartment MLI models in the network to lower computational cost. To preserve the two principal effects conferred by the narrow dendrites of MLIs, namely (i) the passive attenuation of signal predicted by the cable theory, and (ii) the low-pass filtering of somatic membrane potential along the dendrite, the one compartmental models have been amended accordingly. Specifically, the same intrinsic membrane currents included in the ball and stick models are retained in the one compartment MLI models, namely, Na^+^ (*I*_*Na*_), delayed rectifier K^+^ (*I*_*Kdr*_), A-type K^+^ (*I*_*A*_), Ca^2+^-activated K^+^ (*I*_*K*(*Ca*)_), T-type Ca^2+^ (*I*_*T*_ ), HVA Ca^2+^ (*I*_*HVA*_), and leak (*I*_*Leak*_) currents; these are further augmented with additional currents, including hyperpolarization-activated current (*I*_*h*_), and synaptic GABA (*I*_*GABA*_), AMPA (*I*_*AMPA*_), and NMDA (*I*_*NMDA*_) currents. The formalism of each MLI model is given by

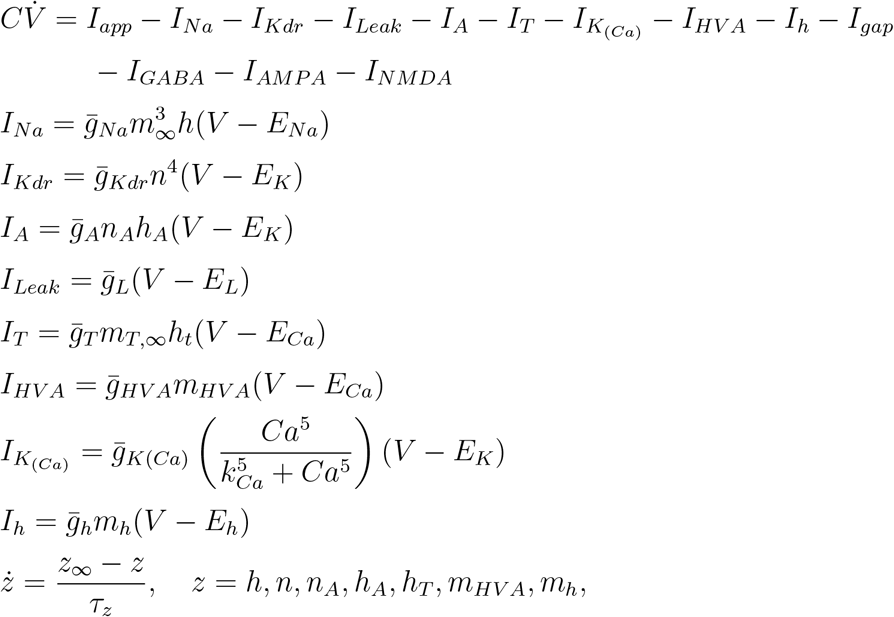

where *I*_*app*_ ∼ 𝒩 (0.0, 0.2) to ensure heterogeneity in firing rate (20 − 55 Hz) across MLIs in line with *in vivo* observations (Brown et al., 2025). Gating properties are unchanged from the ball and stick model (Table 3) and maximal conductances are fit to reproduce *in vitro* firing of CSC (Locatelli et al., 2020) (Figure S2) and are listed in Table 4. The linear non-rectifying gap junctional coupling used for the ball and stick MLI models is updated for neuron *m* (*m* = 1, 2, …, *M* ) and is given by

**Table 4:** Maximal conductances for MLI one compartment model.

| Current | $\bar{g}_x$ (mS/cm<sup>2</sup>) | E (mV) |
| --- | --- | --- |
| Na | 3.0 | 55.0 |
| Kdr | 17.5 | -80.0 |
| A | 7.5 | -80.0 |
| Leak | 0.0125 | -60.0 |
| T | 0.45045 | 22.0 |
| HVA | 0.28 | 22.0 |
| K(Ca) | 0.3 | -80.0 |
| h | 20 | -34.4 |

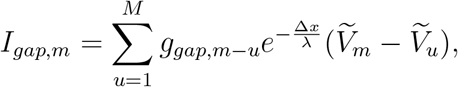

where 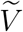 denotes the low-pass filtered *V*, satisfying 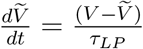, and the factor 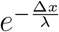 accounts for passive dendritic attenuation based on the intersomatic distance Δ*x* and the cable length constant *λ*, defined earlier from the the ball and stick model. A low-pass filter time constant *τ*_*LP*_ of 28.57 ms is used, corresponding to a a cut-off frequency of 35 Hz.

Granule cell input is approximated as a Poisson point process with a mean frequency of 50 Hz. This reflects the convergence of hundreds of granule cell/parallel fibre synapses onto each MLI (Palay and Chan-Palay, 1974; Carter and Regehr, 2002) as well as the low firing rates of granule cells of about 0.5 Hz *in vivo* (Chadderton et al., 2004). Given that the sum of independent Poisson processes is a Poisson process, with 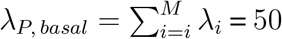 Hz, the excitatory synaptic input from granule cells is approximated as

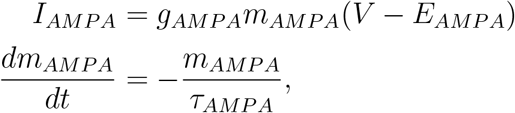

where *E*_*AMPA*_ = 0 mV, *m*_*AMPA*_(*t*) = *m*_*AMPA*_(*t*^−^) + 1 to reflect an increment of 1 at every time point (*t*_GrC_) a granule cell (GrC) fires an action potential, and *τ*_*AMPA*_ = 2 ms is the exponential decay time constant of *m*_*AMPA*_ (Kleppe and Robinson, 1999; Santos-Valencia et al., 2025).

During network simulations, action potentials are detected online using the following criteria: 1. The membrane potential is greater than a threshold *V*_*th*_ = −10 mV (i.e. *V* ≥ *V*_*th*_); 2. The membrane potential of the current time step is smaller than the preceding time step (i.e. *V* (*t*) *< V* (*t*−1)); and 3. No action potential is detected in the preceding 2 ms to ensure that only one single action potential is identified during the down stroke of the spike between its peak and the crossing of the −10 mV threshold. The detected action potentials are then used to increment *m*_*GABA*_ by 1 after a delay of 5.0 ms (Coddington et al., 2013; Lackey et al., 2024) to account for MLI-MLI inhibition (Lackey et al., 2024; Kim and Augustine, 2021; Rieubland et al., 2014; Kim et al., 2014) resulting in a GABAergic synaptic current, given by

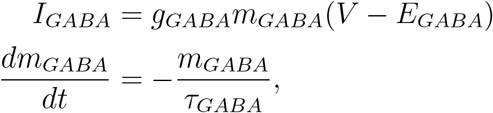

with *E*_*GABA*_ = −80 mV, *m*_*GABA*_(*t*) = *m*_*GABA*_(*t*^−^) + 1, and *τ*_*GABA*_ = 1.9 ms is the exponential decay time constant of *m*_*GABA*_ (Santos-Valencia et al., 2025; Carter and Regehr, 2002).

NMDA receptor activation only occurs at parallel fibre MLI synapses after repetitive or strong stimulation (Carter and Regehr, 2000a; Clark and Cull-Candy, 2002; Nahir and Jahr, 2013). The NMDA current was modelled as a difference of exponentials, but with the addition of the Mg^2+^ block (Wang, 2002; Brunel and Wang, 2001; Jahr and Stevens, 1990), as follows

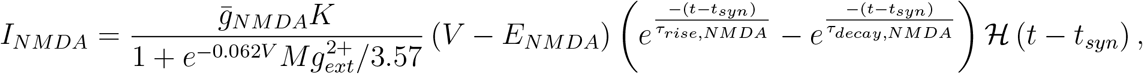

where ℋ is the Heaviside step function, 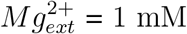 (Wang, 2002; Brunel and Wang, 2001), *τ*_*rise,NMDA*_ = 15 ms, *τ*_*decay,NMDA*_ = 150 ms, and *K* is a normalization factor given by

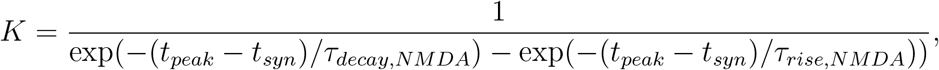

with

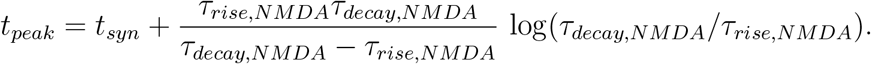

Network connectivity conductances of *g*_*AMPA*_= 1.2 nS, *g*_*GABA*_ = 1.0 nS, *g*_*gap*_ = 0.6 nS (Santos-Valencia et al., 2025) and *g*_*NMDA*_ = 1.8 nS are used unless otherwise stated. Stimulus input consists of transient increases in the rate of the Poisson process for granule cell AMPA input for a duration of 2 ms, the presence of an NMDA current (i.e. *g*_*NMDA*_ *>* 0) or both a Poisson rate increase along with an NMDA current.

### Phase relationships and synchrony quantification

The pairwise phase differences in spike times between 2 cells: *m* and *u*, are computed directly from spike times using the expression

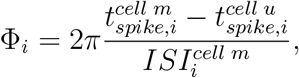

where 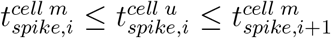 and 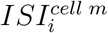 is the interspike interval that follows the spike at time 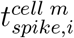. The pairwise phase differences Φ_*i*_ is limited to only capturing 1-to-1 phase locking, as *n*-to-1 locking was not observed in MLI pairs with similar firing rates. An exponential decay is fit to this phase difference between pairs of MLIs to obtain the time constant of synchronization.

Network activity is quantified using the coactivity measure detailed by Brown et al. (2025); it provides a measure of concurrent activity in a network. It is defined as the percentage of neurons in the network with an action potential in a rolling 4 ms window stepped at 1 ms increments. To quantify the synchrony in simulated networks, the mean pairwise phase locking value (PLV; also called mean phase coherence (Mormann et al., 2005)) across all pairs of neurons in the network is calculated over two periods: the baseline period preceding the increase in AMPA input, and for a time window of 50 ms after the increase in basal AMPA input rate. In a pair of neurons, PLV = 1 indicates phase locking, whereas when PLV = 0 indicates lack of synchronization (Tort et al., 2010). For simulations with NMDA input, changes in network synchronization are quantified as the change in mean pairwise PLV (ΔPLV) using the PLV values prior to and in the 50 ms window after the increase in basal Poissonian AMPA input rate. The number of peaks during this 50 ms window with minimum prominence of the greater of 5 times the standard deviation of the coactivity prior to stimulus onset or 1% coactivity are also detected and used as a metric of transient synchronization. Spearman correlations are used to quantify the presence of monotonic relationships between network model parameters and peak coactivity or ΔPLV.

### Simulation Methods

The discretized ball and stick MLI models are simulated using a second order stabilized Runge-Kutta (ROCK2) method (Abdulle and Medovikov, 2001), or an adaptive order quasi-constant timestep numerical differentiation function (QNDF) method (Shampine and Reichelt, 1997) for pairs of models implemented in DifferentialEquations.jl (Rackauckas and Nie, 2017b). Stochastic simulations are performed using the Euler-Maruyama method implemented in StochasticDiffEq.jl (Rackauckas and Nie, 2017a,b). MLI network simulations are performed using adaptive order Adams explicit (VCABM) method (Hairer et al., 1993), implemented in DifferentialEquations.jl (Rackauckas and Nie, 2017b).

### Code accessibility

All code for the modelling are available for download at https://github.com/nkoch1/MLI_synch_2026.git.

## Results

### Ball and Stick MLI model captures firing properties of CSCs

The ball and stick MLI model (Figure 1A) is simulated using a spatial discretization scheme in which the dendritic stick is discretized into *n* = 50 dendritic compartments (see Methods). These simulations are then fit to firing properties of CSCs recorded using whole cell configuration (Locatelli et al., 2020). Specifically, the model is fit to both the action potential phase cycle and the firing-rate responses of CSCs. The resulting model closely reproduces the experimentally observed CSC electrical properties, with the voltage traces (Figure 1B), phase portrait (Figure 1C), and frequency-current curve (Figure 1D), matching those recorded experimentally *in vitro* (Locatelli et al., 2020). Additionally, the model captures somatic action potential backpropagation down the dendritic compartment (Figure 1E), with progressive attenuation in amplitude and increasing delay to their peak as during propagation (Figure 1F).

**Figure 1:**
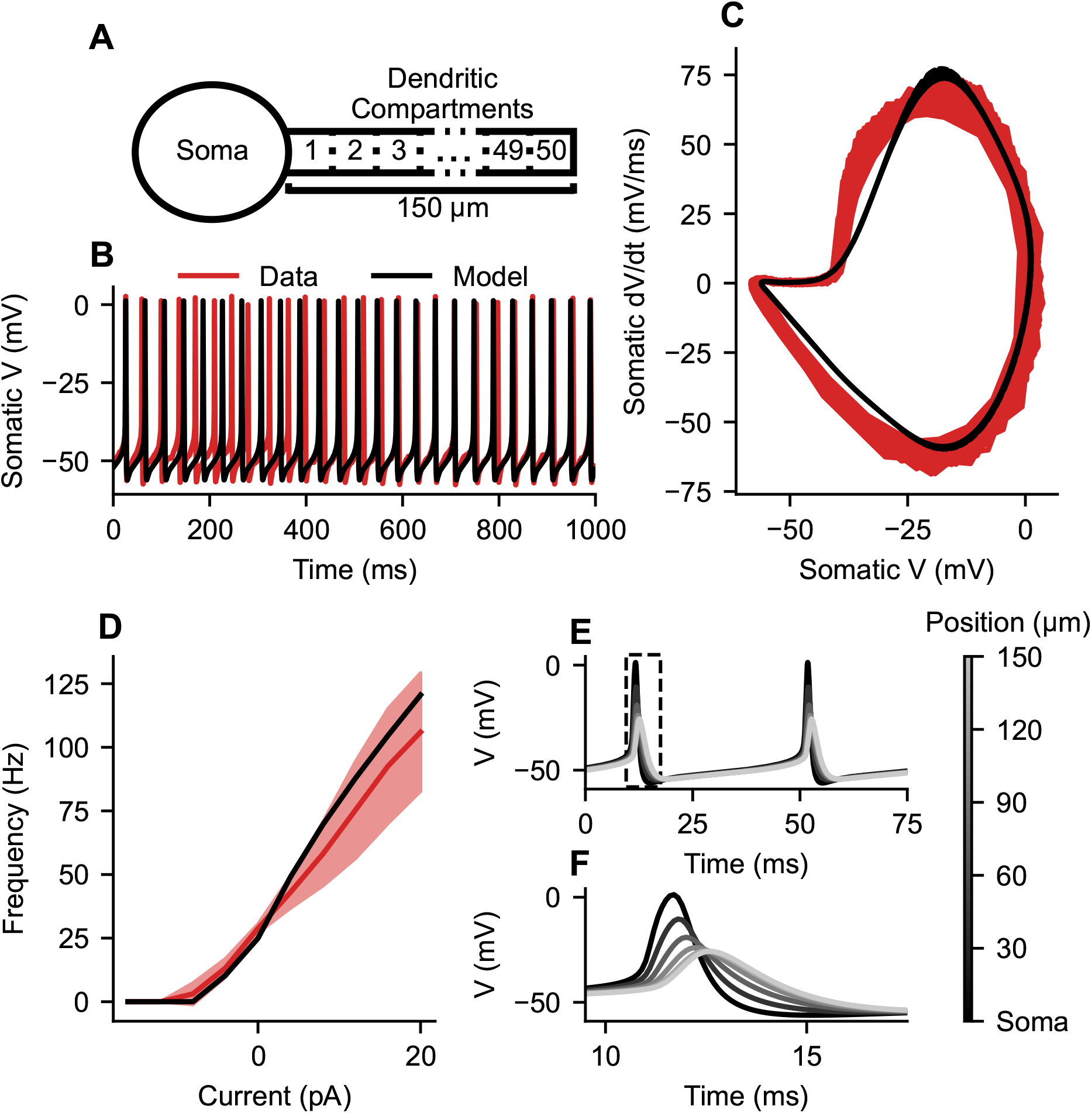
Ball and stick model reproducing MLI firing properties. (A) Schematic depicting the ball and stick model comprised of a somatic compartment and a 150 μm long dendritic cable discretized into 50 compartments. (B) Ball and stick model produces spontaneous somatic membrane potential activity (black), without current injection, resembling MLI activity (red) recorded using gap-free patch clamp protocol (Locatelli et al., 2020). (C) Action potential phase cycles in the (*V, dV/dt*)-plane produced by the MLI ball and stick model (black) and recorded in CSCs (red). (D) Frequency-current curves generated by the MLI ball and stick model (black) and obtained from whole cell patch clamp recordings of CSCs (red) (Locatelli et al., 2020). (E) Action potentials generated in the soma backpropagate along the dendritic cable with attenuation and delay. These action potentials are colour-coded according to the colour-bar, indicating their position relative to the soma. (F) Zoomed-in view of individual action potentials highlighted in the dashed box in E.

### Modest effects of dendritic current removal on dendritic filtering and excitability

To investigate the filtering properties of distal synaptic inputs by the dendrite, an OU noise process is incorporated into the distal dendritic compartment (Figure 2A). Dendritic Kdr K^+^ (Figure 2B), A-type K^+^ (Figure 2C), T-type Ca^2+^ (Figure 2D), HVA Ca^2+^ (Figure 2E) and K(Ca) K^+^ (Figure 2F) currents are then individually removed to investigate their role in the filtering of the OU noise representing synaptic bombardment. When the gain across the distal dendritic compartment and somatic membrane potentials is considered (Figure 2G), minor change in the frequency dependence of the dendritic filtering is observed following the removal of dendritic currents, suggesting that the passive dendritic properties dictate filtering of synaptic inputs.

**Figure 2:**
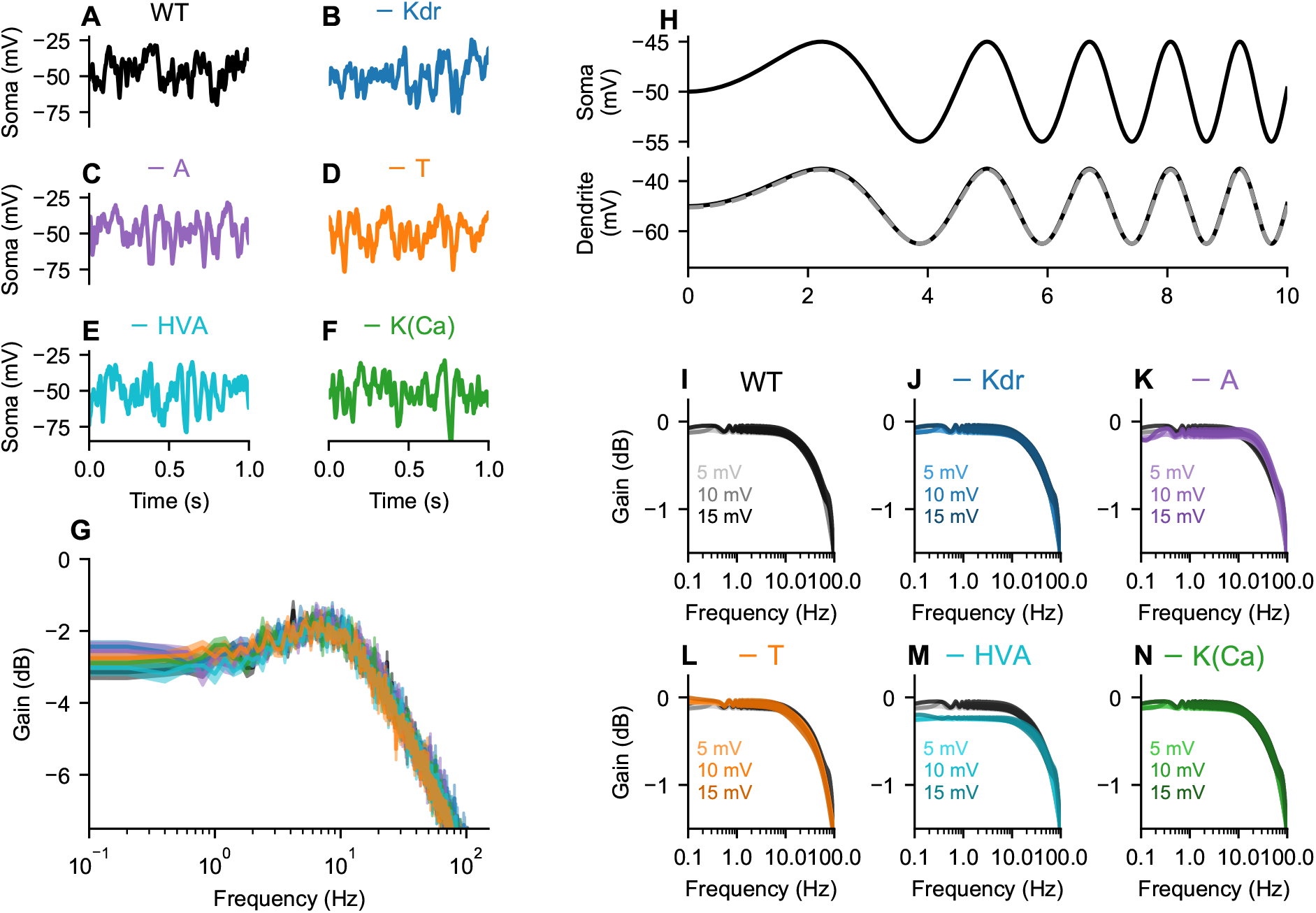
Filtering properties of the dendritic cable and the role of dendritic currents. (A-F) Effect of Ornstein-Uhlenbeck noise (*θ* = 0.5 ms^−1^, *μ* = *E*_*Leak*_, *σ* = 30) injected into the distal dendritic compartment of the MLI ball and stick model on somatic membrane potential in the original model (A), and in models in which individual dendritic currents are removed, including delayed rectifier K^+^ (B; Kdr), A-type K^+^ (C; − A), T-type Ca^2+^ (D; − T), high voltage activated Ca^2+^ (E; − HVA) and Ca^2+^ activated K^+^ (F; − K(Ca)) currents. (G) The resulting gain between the input Ornstein-Uhlenbeck noise and the somatic response. (J) chirp stimulus consisting of a sinusoidal input with linearly increasing frequency injected into the soma (top). The stimulus spans a frequency range of (0.001 - 100 Hz and is applied at three amplitudes 5, 10 and 15 mV over 1000 s). Example dendritic responses (bottom) for the most proximal (black) and most distal (grey) dendritic compartments in response to the chirp stimulus. (I-N) The gain across the most distal and most proximal compartments calculated for three chirps of 5, 10 and 15 mV amplitudes injected into the some of the original model (I), and in models in which individual dendritic currents are removed, including delayed rectifier K^+^ (J; − Kdr), A-type K^+^ (K; − A), T-type Ca^2+^ (L; − T), high voltage activated Ca^2+^ (M; − HVA) and Ca^2+^ activated K^+^ (N; − K(Ca)).

However, the backpropagation of action potentials from the soma along the dendrite still occurs (Figure 1E-F). To investigate the filtering of subthreshold somatic membrane potential fluctuations during dendritic backpropagation, a chirp stimulus consisting of a sinusoidal somatic membrane potential that increases frequency linearly over time at 3 amplitudes (5, 10 and 15 mV (Figure 2H top) is applied. The distal and proximal dendritic membrane potentials (grey and black, respectively; Figure 2H bottom) are then used to compute the dendritic gain of the chirp (Figure 2I). Removal of dendritic Kdr K^+^ (Figure 2J), A-type K^+^(Figure 2K), T-type Ca^2+^ (Figure 2L), HVA Ca^2+^ (Figure 2M) and K(Ca) K^+^ (Figure 2N) currents do not generally change the frequency dependency profiles of the filtering. However, removal of A-type K^+^ current reduces (increases) attenuation of higher (lower) frequencies above 10 Hz (below 1 Hz) compared to the original model (Figure 2K). Removal of dendritic HVA Ca^2+^ current, on the other hand, is limited to the attenuation of frequencies below 15 Hz (Figure 2M). Overall, the limited effects of dendritic current removal on the filtering of somatic membrane potential chirp indicates that subthreshold somatic oscillations are filtered predominantly as a result of the passive properties of the dendrites, in agreement with the observations obtained using distal OU input.

It is important to note that although the input-output function (i.e. frequency-current curve) of the cable model shifts slightly with the removal of individual dendritic currents (especially HVA, T, A and Kdr), the overall frequency-current curve shape is largely maintained (Figure S3), indicating that the individual removal of these dendritic currents is not fundamentally altering the type of excitability of the model, but are rather modestly shifting the firing frequency.

### Gap junctional strength, location, and initial phase govern MLI synchronization

To examine how gap junctional coupled affects the synchrony of MLIs, pairs of ball and stick MLI models are coupled together through dendro-dendritic linear gap junctions. Such gap junctions can result in the synchronization of firing in a MLI pair (Figure 3A) with the phase difference (ΔΦ) converging over time to a steady state ΔΦ_*ss*_ (Figure 3B) as illustrated by the concentric circular time plot with time extending outward. To examine the effect of gap junctional coupling on synchronization, the initial ΔΦ between MLI pairs (Figure 3C), the strength of the gap junctional conductance (*g*_*gap*_; Figure 3D), and the distance of the gap junction from the soma (Figure 3E) are altered. As the initial ΔΦ increases from 0 or 2*π*, the time to synchronize to a steady state ΔΦ_*ss*_ increases with the slowest synchronization occurring with an anti-phase initialization (*π*; Figure 3C). A pair of MLIs can synchronize with either neuron leading in phase, resulting in steady state phase differences ΔΦ_*ss*_ that could be positive or negative (Figure 3C). Alteration of the magnitude of the gap junctional conductance at a fixed initial phase difference: ΔΦ = 1*/*3*π*, reveals that a stronger coupling promotes faster synchronization, whereas intermediate coupling strength produces a non-zero steady state ΔΦ_*ss*_ (Figure 3D). Consistently, increasing the distance of gap junctional coupling from the soma decreases the coupling strength and results in smaller changes in ΔΦ_*ss*_ from the initial ΔΦ (Figure 3E).

**Figure 3:**
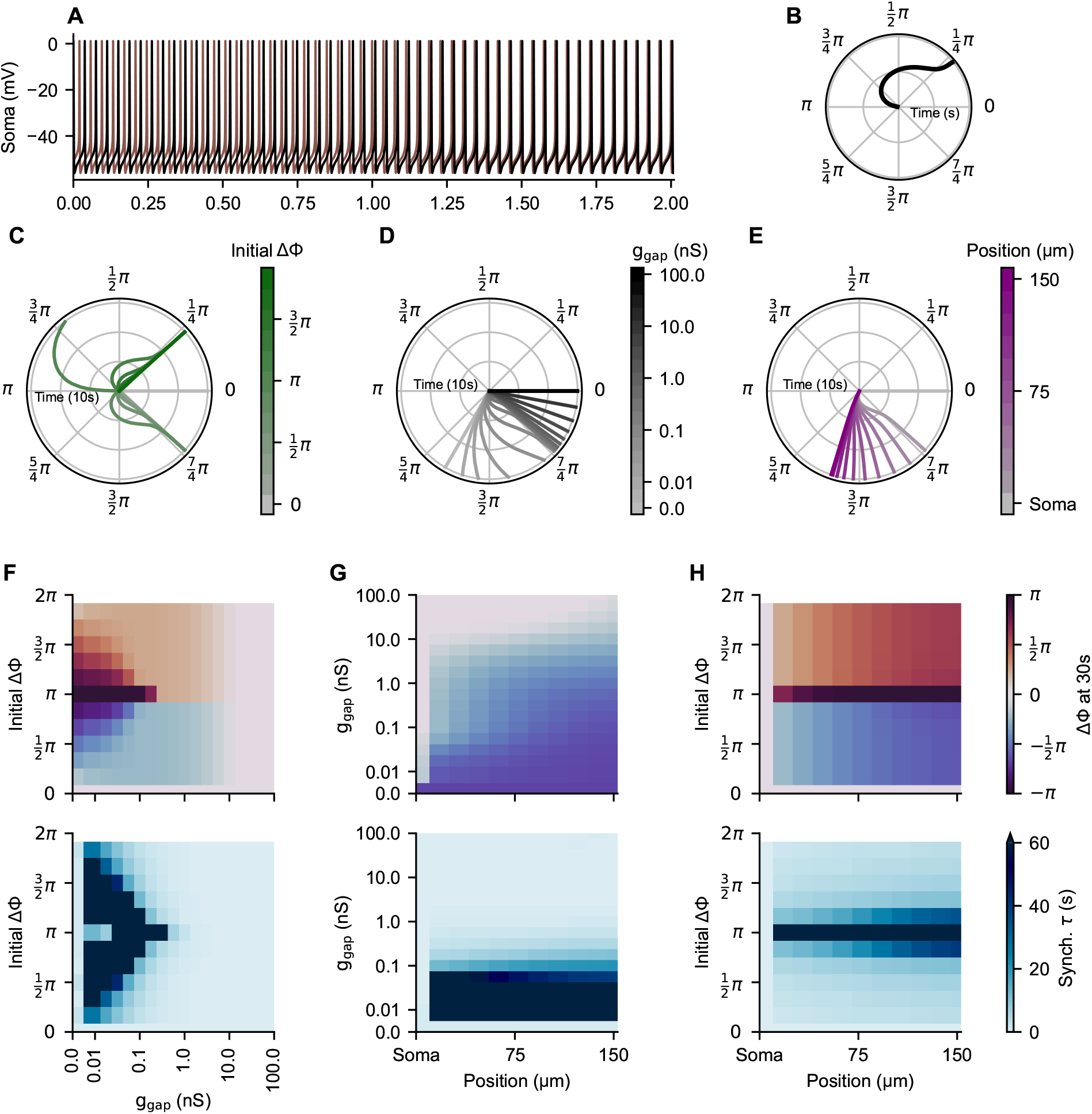
Effects of gap junctional position and conductance, and initial phase difference on MLI synchronization. (A) Simulated spontaneous firing activity of a pair of ball and stick models (black and brown) coupled with a gap junction (30 μm from soma with gap junctional conductance*g*_*gap*_ = 2.0 nS and initial phase different ΔΦ = *π*), showing synchronized activity over time . (B) Concentric circular time plot quantifying the time evolution of the phase difference between the action potentials of the two models in (A). Time axis starts in the centre of each plot and expands outward with each circle indicating the passage of 10 s. Note that phase difference magnitude is dependent on which model is the reference. (C-E) Concentric circular time plot of a gap junctionally coupled MLI pair depends on the initial phase difference (C; gap junction 15 μm from soma with *g*_*gap*_ = 0.178 nS), gap junctional conductance (D; gap junction 15 μm from soma with an initial ΔΦ = *π/*3) and position of the gap junction along the dendrite (E; *g*_*gap*_ = 0.178 nS and initial ΔΦ = *π/*3). (G-H) Heatmaps showing the effect of (F) initial ΔΦ and *g*_*gap*_ (gap junction 15 μm from soma), (G) *g*_*gap*_ and gap junctional position (initial ΔΦ = *π/*3), and (H) junctional position and initial ΔΦ (*g*_*gap*_ = 0.178 nS) on the phase difference after 30 s (ΔΦ_30_; top) and the synchronization time constant (*τ* ) computed by fitting an exponential decay to the phase differences for the simulations in F-H. Heatmaps are colour-coded according to colour bars on the right.

To further examine how these three factors influence synchronization in pairs of gap junctionally coupled ball and stick MLI models, a systematic characterization of their effects is performed. This is initially done by investigating the interaction between initial ΔΦ and *g*_*gap*_ on both ΔΦ_*ss*_ (Figure 3F top) and synchronization time (Figure 3F bottom). Our results reveal that, at intermediate *g*_*gap*_, a non-zero ΔΦ_*ss*_ is observed with synchronization time decreasing as *g*_*gap*_ increases to produce stronger coupling. At high *g*_*gap*_, complete synchronization occurs with short synchronization timescale across all initial ΔΦ (Figure 3F); in contrast, at low *g*_*gap*_, complete synchronization is not achieved due to the very slow timescale of convergence, exceeding 30 s of simulated time (Figure 3F). Expanding this analysis to examining the combinatorial effects of *g*_*gap*_ and gap junctional position along the dendrite on ΔΦ_*ss*_ (Figure 3G top) and timescale of synchronization (Figure 3G bottom) suggest that at intermediate *g*_*gap*_, ΔΦ_*ss*_ depends on gap junctional strength and distance from the soma with small conductance gap junctions resulting in slow synchronization with time constants greater than 1 min (Figure 3G). To further explore the distance dependent effect, the impact of gap junctional position is considered in combination with the initial ΔΦ (Figure 3H). In this case, ΔΦ_*ss*_ increases with distance across initial ΔΦ (Figure 3H top) and the timescale of synchronization increases with distance at initial ΔΦ near *π* (Figure 3H bottom). Taken together, these results indicate that smaller (larger) conductance more distal (proximal) dendro-dendritic coupling from the soma are less (more) effective at promoting rapid synchronization between pairs of MLIs, in line with previous findings in other neurons (Pfeuty et al., 2005, 2007; Saraga and Skinner, 2004; Remme et al., 2012; Schwemmer and Lewis, 2014; Vervaeke et al., 2010; Mendoza and Haas, 2022).

### Modest effects of dendritic current removal on MLI synchronization

Although the effect of dendritic currents in the ball and stick models on dendritic filtering is minimal (Figure 2), these currents may nonetheless contribute to synchronization in a manner not captured by the the dendritic filtering investigation. To address this, the impact of dendritic currents on synchronization of gap junctionally coupled MLI pairs of ball and stick models is assessed by the removal of dendritic currents one at a time across a range of gap junctional conductances and locations along the dendrite for a sequence of 12 initial ΔΦ. This is accomplished by quantifying both ΔΦ at 30 s (ΔΦ_30_) and synchronization time constant as a function of *g*_*gap*_ and the dendro-dendritic coupling location relative to the soma, with results summarized as heatmaps for each initial ΔΦ. In the case of the original model, ΔΦ_30_ and synchronization time constant (Figure 4A top and bottom, respectively), generally exhibit effects consistent with our previous results (Figure 3G), showing that across initial ΔΦ with increasing *g*_*gap*_ and more proximal gap junctions result in greater coupling and synchrony. This feature persists even when dendritic Kdr K^+^ (Figure 4B), A-type K^+^ (Figure 4C), T-type Ca^2+^ (Figure 4D), HVA Ca^2+^ (Figure 4E) and K(Ca) K^+^ (Figure 4F) currents are individually removed. Notably, when the pair of MLI models is initialized at ΔΦ = 0, we obtain synchrony in all conditions, whereas when the pair of MLI models is initialized at a non-zero initial ΔΦ, rapid synchronization at 30 s is achieved only when *g*_*gap*_ is sufficiently large or the gap junction is in close proximity to the soma.

**Figure 4:**
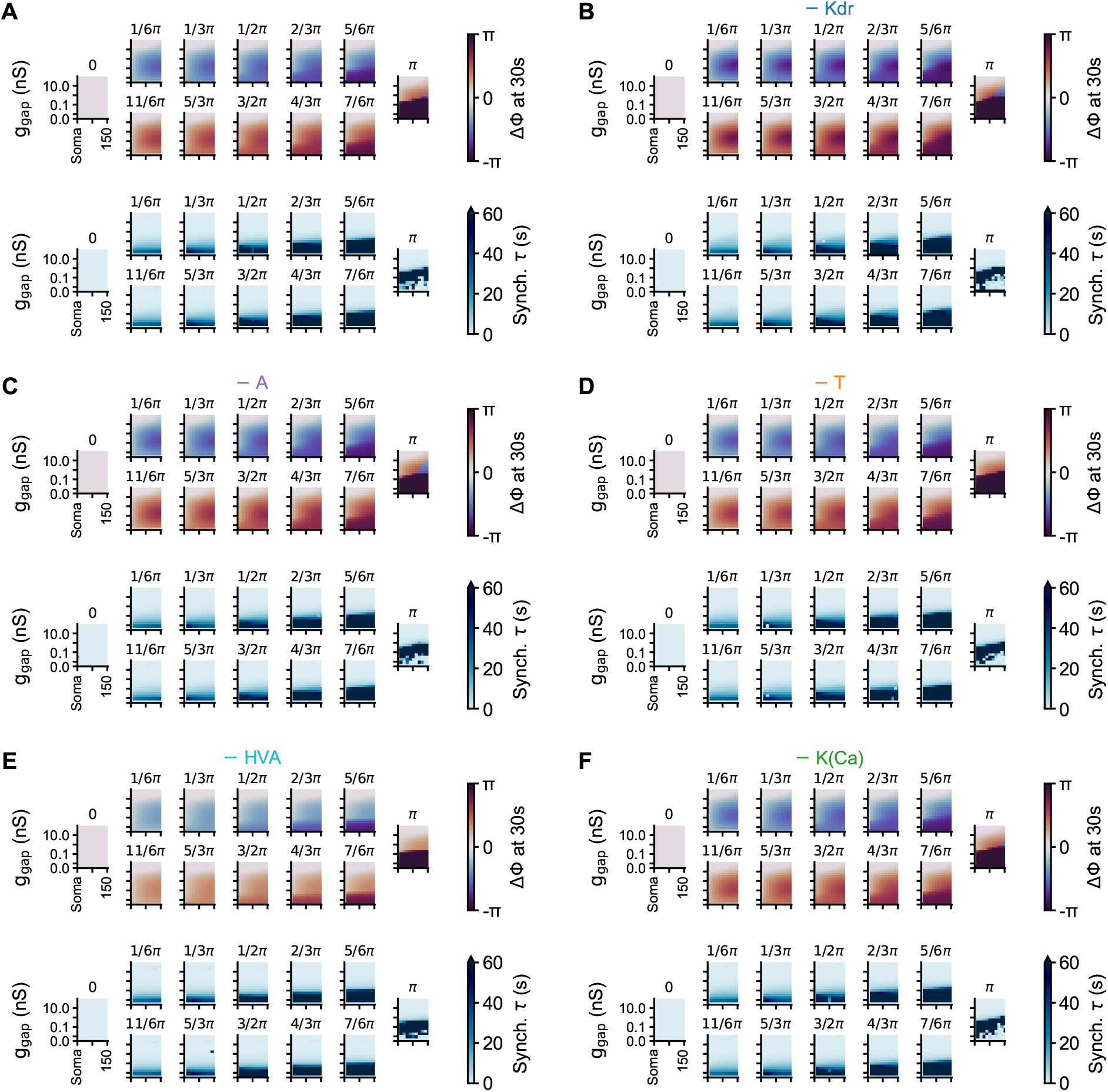
Synchronization in gap junctionally coupled MLI pairs during dendritic current removal. (A) The phase difference at 30 s (ΔΦ_30_, top) and synchronization time constant (bottom) obtained by exponential fitting of the phase difference (Synch. *τ*, bottom) for a pair of gap junctionally coupled original MLI models across gap junctional position (x-axis) and gap junction conductance (*g*_*gap*_; y-axis) for initial phase differences indicated on top of each panel (top row initial phase differences are equivalent to the bottom row with opposite reference model). (B-F) The same as in (A) with individual dendritic currents removed, including delayed rectifier K^+^ (B; − Kdr), A-type K^+^ (C; − A), T-type Ca^2+^ (D; − T), high voltage activated Ca^2+^ (E; − HVA) and Ca^2+^ activated K^+^ (F; − K(Ca)) currents.

It is important to note, however, that although the removal of dendritic Kdr K^+^ (Figure 4B), A-type K^+^ (Figure 4C), T-type Ca^2+^ (Figure 4D), HVA Ca^2+^ (Figure 4E) and K(Ca) K^+^ (Figure 4F) currents does not generally alter the phase differences of the gap junctionally coupled MLI or the synchronization timescale compared to the original model (Figure 4A), modest deviations are observed under specific conditions: removal of dendritic HVA Ca^2+^ current moderately decreases ΔΦ_30_ at intermediate *g*_*gap*_ (Figure 4E), whereas removal of dendritic Kdr K^+^ currents increases ΔΦ_30_ at intermediate *g*_*gap*_ and distal gap junctions (Figure 4B). The ΔΦ_30_ and synchronization timescale increases with increasing initial ΔΦ.

Collectively, these results suggest that synchronization remains most strongly governed by gap junctional properties, with low *g*_*gap*_ resulting in longer synchronization times and correspondingly larger ΔΦ_30_ between pairs of MLI models. Indeed, across all dendritic current manipulations, the qualitative synchronization behaviour is preserved, indicating that MLI coupling dynamics are primarily determined by *g*_*gap*_ and the spatial localization of the gap junction relative to the soma.

### GABAergic inhibition modulates AMPA-driven peak synchronization of MLI network

MLIs are gap junctionally coupled to more than one other MLI (Kim et al., 2014; Mann-Metzer and Yarom, 1999; Alcami and Marty, 2013); therefore, examining synchronization in pairs of MLI models, while informative about the role of intrinsic MLI properties on gap junctional synchronization, is likely not reflective of the electrical coupling present in MLI networks. To investigate how gap junctional coupling contributes to MLI network activity, 100 network realizations of *M* = 336 one compartment MLI models (rather than ball and stick models to reduce computational complexity; see Methods) are created, with each MLI receiving a Poissonian granule cell AMPA input and being coupled to a variable number of other MLIs via probabilistic gap junctions and inhibitory synapses (see Methods). These network models are then simulated to investigate the underlying components responsible for generating the activity of MLIs recorded *in vivo* in response to a sensory stimulus (Brown et al., 2025).

We initially assess whether the stimulus-evoked MLI firing increases recorded *in vivo* is the result of AMPA input by transiently increasing the rate of the Poissonian AMPA input into each MLI model in the network for 2 ms. Representative firing activity from a single network realization, together with coactivity traces from 100 realizations of the network model are then quantified in response to four-fold (Figure 5A), ten-fold (Figure 5B), and sixteen-fold (Figure 5C) increases in the basal AMPA input rate (*λ*_*P,basal*_) applied at 0 ms; in all cases, this transient increase in basal AMPA input rate elicits a marked rise in firing in many MLIs in the network and increases the coactivity in the network briefly in a manner dependent on the magnitude of the input rate change (Figure 5A-C). When a broader range of basal AMPA input rate changes is considered, the peak coactivity across the networks rises monotonically with larger AMPA drive, reaching values approaching 80% for the largest input elevations (Figure 5D; Spearman *ρ* = 0.989, p ≤ 0.001). Since an increase of fourteen-fold in the basal AMPA input rate (*λ*_*P,basal*_) is sufficient to elicit a response similar in magnitude to the largest peak coactivity observed by Brown et al. (2025) with highest peak coactivity, this fold increase in *λ*_*P,basal*_ is used hereafter (unless otherwise stated).

**Figure 5:**
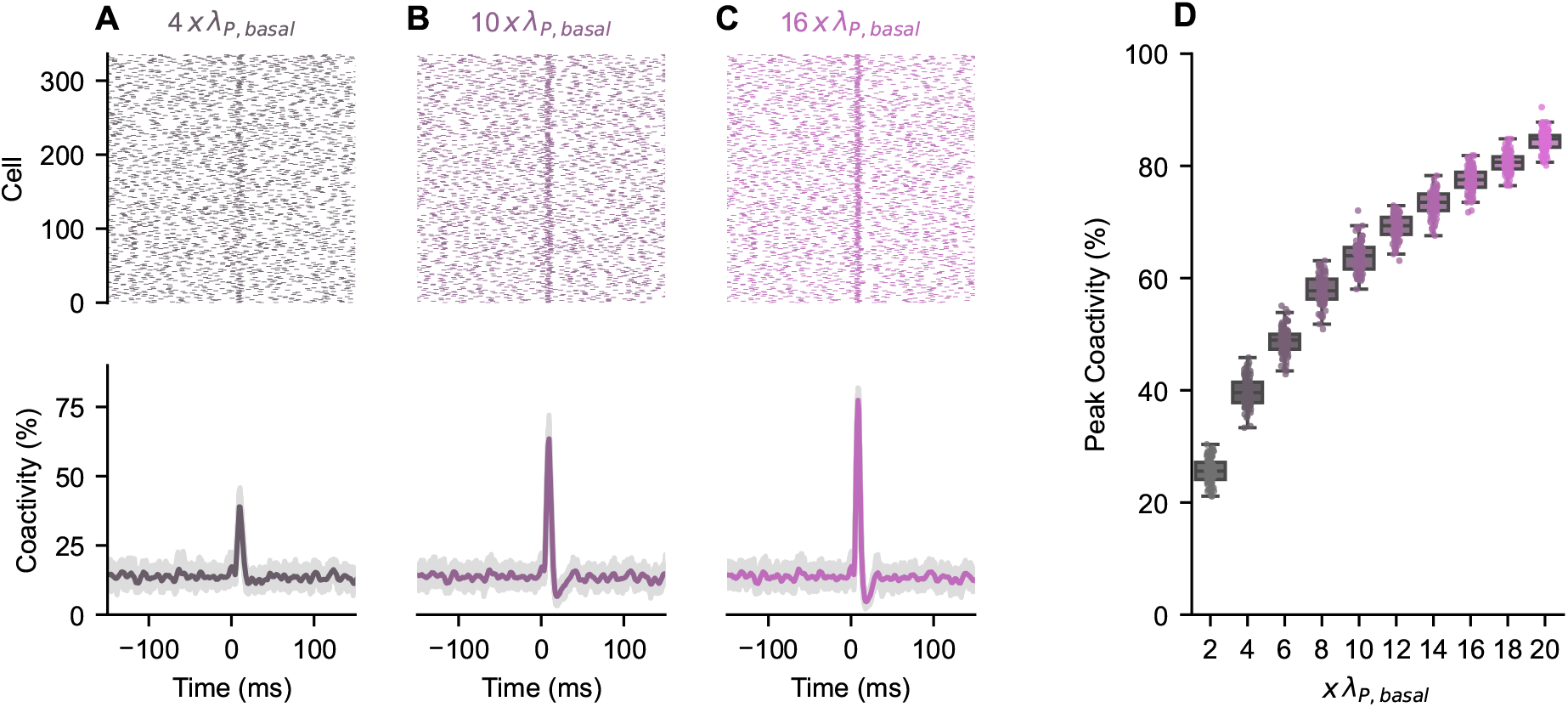
Effect of increasing AMPA input transiently on MLI network synchronization. (A-C) Raster plots from the same network realization (top), together with coactivity traces across *n* = 100 network realizations (in light gray; bottom) and their mean coactivity (in colour; bottom), showcasing the effect of transiently increasing the rate of the Poisson point process representing granule cell AMPA input onto MLIs. AMPA basal rate ( *λ*_*P, basal*_) is increased four-fold (A), ten-fold (B) and sixteen-fold (C) at time 0 ms for 2 ms for the entire network of MLI models. (D) Peak coactivity across all *n* = 100 network realizations when increasing the rate of granule cell AMPA input (x *λ*_*P,basal*_).

Next, the role of gap junctional and synaptic connections between MLIs within the network models in shaping evoked responses to transient increases in AMPA input rate (*λ*_*P,basal*_) is assessed. Increasing the strength of gap junctional coupling (*g*_*gap*_) in the MLI network from 0.0 nS (Figure 6A) to 1.2 nS ((Figure 6), and to 3.0 nS (Figure 6C) produces only modest changes in peak coactivity. This trend generally holds across 100 network realizations (Figure 6D) over the full range of *g*_*gap*_ values considered, although larger *g*_*gap*_ is associated with a small increase in peak coactivity (Spearman *ρ* = 0.770, p ≤ 0.001). In contrast, increasing the MLI-MLI inhibition in the network by increasing *g*_*GABA*_ from 0.0 nS (Figure 6E) to 2.0 nS (Figure 6F), and then to 5.0 nS (Figure 6G) decreases the peak coactivity evoked by the transient increase in *λ*_*P,basal*_ and occurs generally as a result of increased *g*_*GABA*_ (Spearman *ρ* = −0.907, p ≤ 0.001) in the 100 realization of the network (Figure 6H). The greater modulation of peak coactivity in response to the 2 ms AMPA input by GABAergic inhibition compared to gap junctional coupling occurs across different *g*_*gap*_ and *g*_*GABA*_ (Figure 6I). This suggests that the peak coactivity of MLIs seen *in vivo* in response to sensory stimuli (Brown et al., 2025) likely reflect a transient increase in AMPA input to MLIs, with MLI-MLI GABAergic inhibition playing a prominent role while gap junctional coupling contributing modestly to modulating MLI responses to this input. That said, the elevation in firing and damped oscillations in coactivity seen after large peak coactivity responses seen *in vivo* (Brown et al., 2025) cannot, however, be attributed to MLI network response to such an exclusively AMPA input.

**Figure 6:**
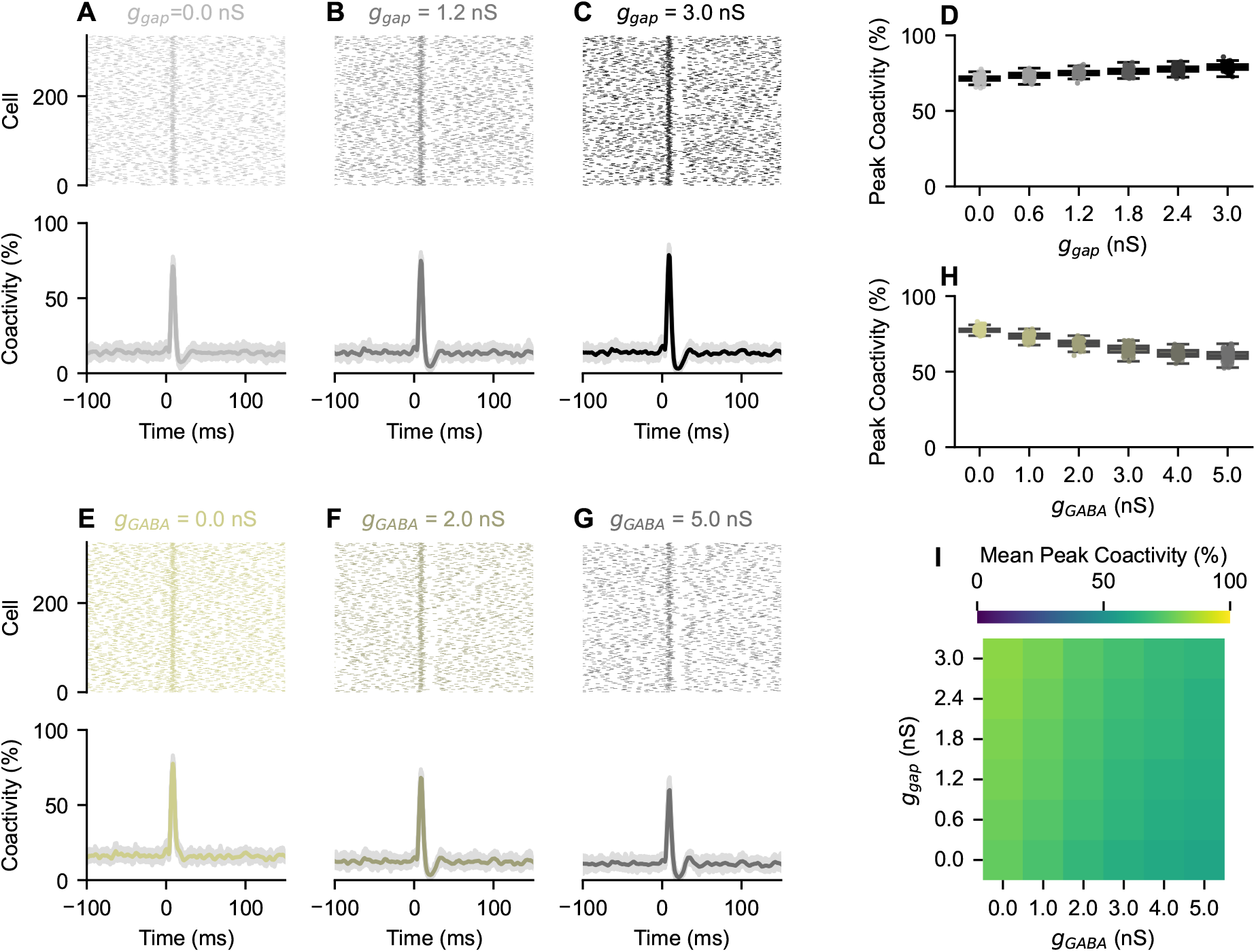
Effect of gap junctional and GABAergic conductances on MLI network synchronization. (A-C) Raster plots from the same network realization (top), together with coactivity traces across *n* = 100 network realizations (in light gray; bottom) and their mean coactivity (in colour; bottom), showcasing the effect of altering gap junctional conductance (*g*_*gap*_) on MLI network synchrony. Network activity is generated by setting *g*_*gap*_ to 0 (A), 1.2 (B), and 3.0 (C) nS in response to a fourteen-fold transient increase in granule cell AMPA input at time 0 ms for 2 ms. (D) Peak coactivity across *n* = 100 network realizations as a function of *g*_*gap*_. (E-F) The same as in (A-C) except that *g*_*gap*_ is replaced by MLI-MLI inhibitory conductance (*g*_*GABA*_). Network activity is generated by setting *g*_*GABA*_ to 0 (E), 2 (F), and 5 (G) nS in response to a fourteen-fold transient increase in granule cell AMPA input at time 0 ms for 2 ms. (H) The same as in (D) except that *g*_*gap*_ is replaced by *g*_*GABA*_. (I) Heatmap showing the effect of altering both *g*_*gap*_ and *g*_*GABA*_ on mean peak coactivity. Heatmap is colour-coded according to the colour-bar on top.

### NMDA input and gap junctional coupling promote prolonged elevated firing and transient MLI network synchronization

If glutamatergic synaptic input is sufficiently large, glutamate spillover can activate extrasynaptic NMDA receptors in MLIs (Nahir and Jahr, 2013; Carter and Regehr, 2000b; Clark and Cull-Candy, 2002; Szapiro and Barbour, 2007; Jörntell and Ekerot, 2002, 2003; Kim and Augustine, 2021), generating an NMDA-mediated current in these cells. To reflect this in our study, an NMDA current is added to the MLI models in the network (see Methods) to test the hypothesis that NMDA activation results in the elevated firing and damped oscillations of coactivity observed in Brown et al. (2025).

The inclusion of an NMDA current with *g*_*NMDA*_ = 1.2 nS produces a prolonged elevation in firing after a four-fold increase in AMPA input basal rate (*λ*_*P,basal*_; Figure 7A), in line with the prolonged increase in firing seen *in vivo* (Brown et al., 2025). Increasing *λ*_*P,basal*_ transiently twenty-fold its basal rate results in larger synchronized AMPA responses followed by damped oscillatory activity leading to similar elevated *in vivo* firing activity (Figure 7B). Increasing AMPA input, while keeping NMDA input constant (*g*_*NMDA*_ = 1.2 nS), during the combined AMPA and NMDA input, generally increases the peak coactivity (Figure 7C; Spearman *ρ* = 0.977, p ≤ 0.001) in a manner similar to the effect seen when AMPA input is applied exclusively. Specifically, increases in *λ*_*P,basal*_ does not meaningfully alter the synchronization in the 50 ms post-stimulus measured by the change in mean pairwise PLV during this window compared to before stimulus application (ΔPLV) (Figure 7D; Spearman *ρ* = 0.271, p ≤ 0.001), suggesting that AMPA drives the initial brief large network response but does not affect the synchronization and elevated firing afterwards.

**Figure 7:**
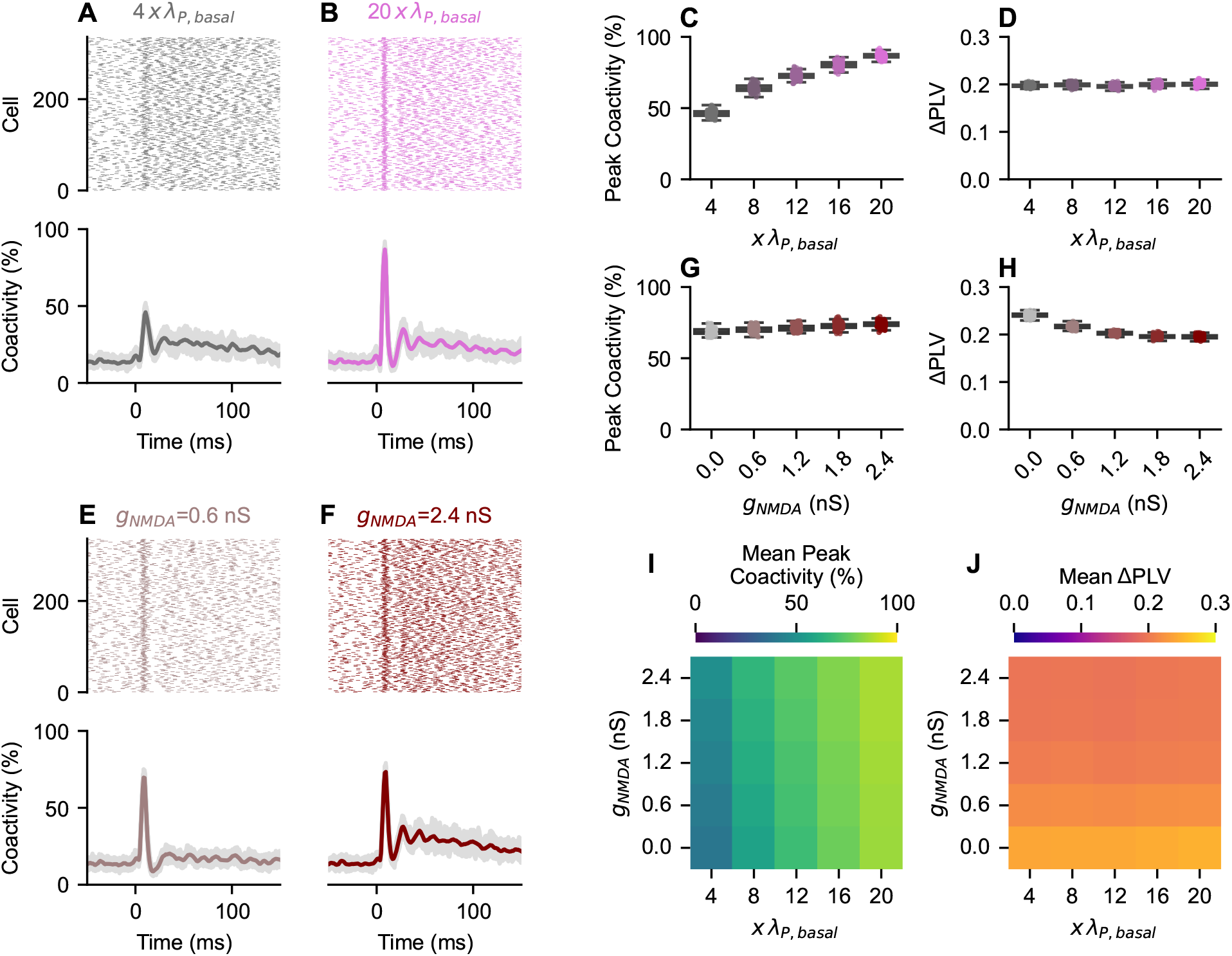
Combined effect of AMPA and NMDA inputs on peak coactivity and pairwise phase locking in the MLI networks. (A,B) Raster plots from the same network realization (top), together with coactivity traces across *n* = 100 network realizations (in light gray; bottom) and their mean coactivity (in colour; bottom), showcasing the effect of altering basal AMPA input (*λ*_*P, basal*_) on MLI network synchrony. Network activity is generated by increasing basal granule cell AMPA input rate four-fold (A), and twenty-fold (B) its basal level *λ*_*P, basal*_ at time 0 ms for 2 ms. (C,D) Summary of peak coactivity (C) and change in mean pairwise phase locking value (ΔPLV) (D) across *n* = 100 network realizations as a function of fold change in *λ*_*P, basal*_. (E,F) The same as in (A,B) except that *λ*_*P, basal*_ is replaced by the NMDA input rate (*g*_*NMDA*_). Network activity is generated by setting *g*_*NMDA*_ to 0 (E), and 2.4 (F) nS in response to a fourteen-fold transient increase in granule cell AMPA input at time 0 ms for 2 ms. (G,H) The same as in (C,D) except that *λ*_*P, basal*_ is replaced by *g*_*NMDA*_. (I,J) Heatmaps showing the effect of altering both *λ*_*P, basal*_ and *g*_*NMDA*_ on mean peak coactivity (I) and mean (ΔPLV) (J). Heatmaps are colour-coded according to colour-bars on top of each panel.

This has prompted us to investigate the role of NMDA input in generating the prolonged elevated response. Consistent with the hypothesis that this sustained activity is NMDA-dependent, small amplitude NMDA input (*g*_*NMDA*_ = 0.6 nS) results in modest elevation of firing rates (Figure 7E), whereas larger NMDA input produces increased firing (Figure 7F), suggesting that the elevated firing is indeed due to NMDA input. Interestingly, increasing the NMDA component of the input (with constant fourteen-fold increase in *λ*_*P,basal*_) also induces small increases in the peak coactivity (Figure 7G; Spearman *ρ* = 0.661, p ≤ 0.001). However, the increase in NMDA amplitude during fixed AMPA input decreases ΔPLV (Figure 7H; Spearman *ρ* = −0.910, p ≤ 0.001) especially as *g*_*NMDA*_ increases from 0.0 to 2.4 nS. This is likely due to the effects of increased firing rate driven by increased NMDA input, leading to a partial desynchronization of the AMPA-evoked transient synchrony. That is, increased NMDA input during afterhyperpolarization together with heterogeneity in connectivity and firing rate results in less synchronicity in the next action potential provided that this general depolarizing input is more dominant than the gap junctional coupling effect. In line with this, removal of gap junctions from the network (i.e. by setting *g*_*gap*_ = 0) results in a similar decrease in ΔPLV with increasing *g*_*NMDA*_ (Figure S4), suggesting that this decrease is independent from the electrical coupling of MLIs in the network. Collectively, these results suggest that transient NMDA and gap junction dependent MLI synchronization following AMPA input is most prominent at large AMPA and modest NMDA inputs (Figure 7I-J).

### Gap junctional coupling and GABAergic inhibition shape NMDA-mediated MLI network synchronization

The effect of modulating gap junctional connectivity on network behaviour is further assessed by altering the gap junctional conductance (*g*_*gap*_) across the network. Setting *g*_*gap*_ = 0 nS causes a loss of oscillatory firing in coactivity at the population level despite exhibiting elevated firing rate post AMPA driven response (Figure 8A); this is in direct contrast to the damped oscillatory population coactivity seen when setting (*g*_*g*_*ap* = 3 nS; Figure 8B). Interestingly, increases in gap junctional conductance is associated with a small increase in peak coactivity (Figure 8C; Spearman *ρ* = 0.750, p ≤ 0.001) similar to those observed when only changing the AMPA input rate (Figure 6D). However, synchronization as measured by ΔPLV in the 50 ms post-stimulus increases as the *g*_*gap*_ increases (Figure 8D; Spearman *ρ* = 0.860, p ≤ 0.001) especially at low *g*_*gap*_. It follows that gap junctional coupling directly contributes to the damped oscillatory population level coactivity seen during elevated response *in vivo* (Brown et al., 2025) and is dependent on the slow depolarization evoked by NMDA receptor activation.

**Figure 8:**
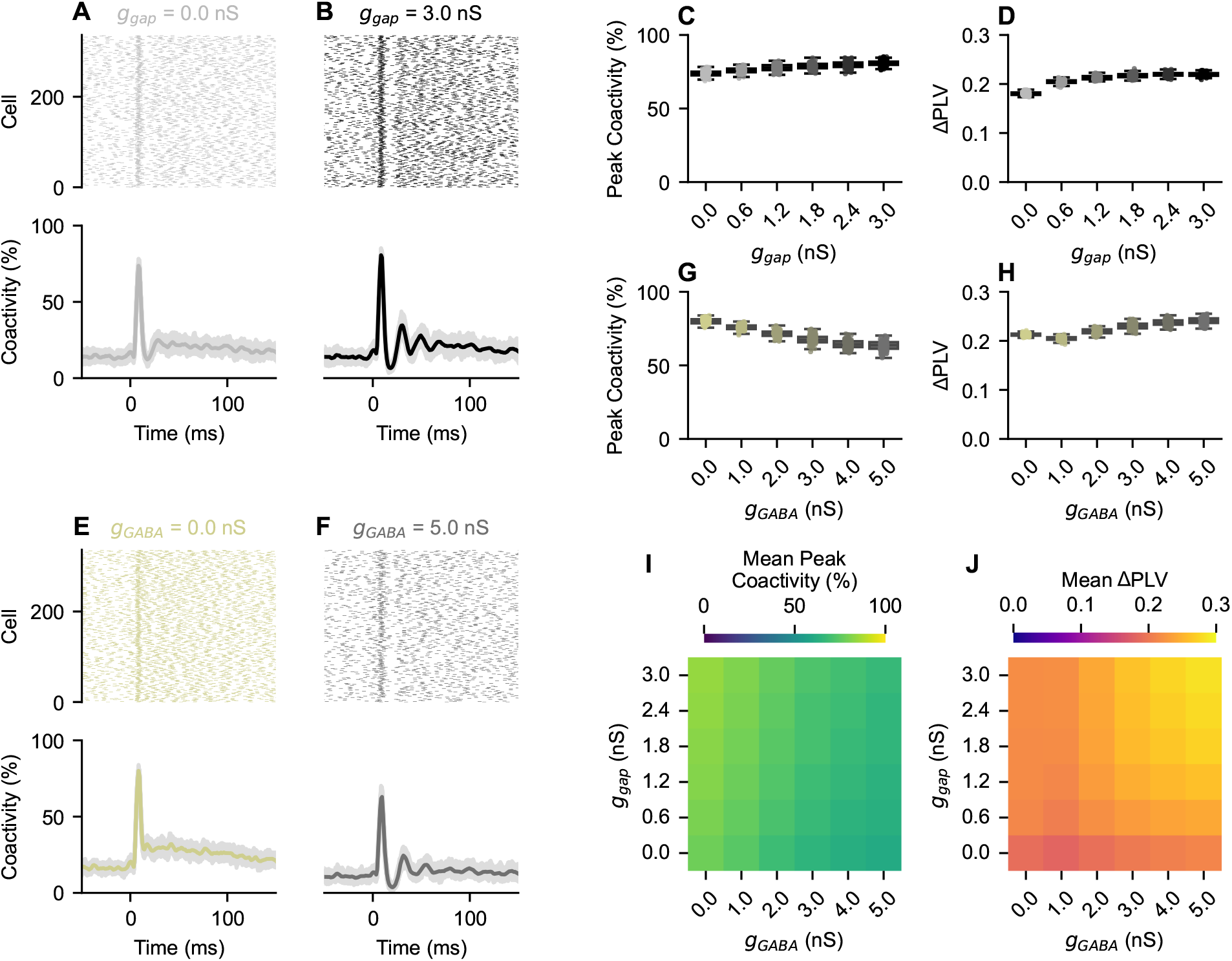
Combined effect of gap junctional and GABAergic conductances on peak coactivity and pairwise phase locking in the MLI networks. (A,B) Raster plots from the same network realization (top), together with coactivity traces across *n* = 100 network realizations (in light gray; bottom) and their mean coactivity (in colour; bottom), showcasing the effect of altering gap junctional conductance (*g*_*gap*_) on MLI network synchrony. Network activity is generated by setting *g*_*gap*_ to 0 (A), and 3 (B) nS in response to a fourteen-fold transient increase in granule cell AMPA input at time 0 ms for 2 ms. (C,D) Summary of peak coactivity (C) and change in mean pairwise phase locking value (ΔPLV) (D) across *n* = 100 network realizations as a function of *g*_*gap*_. (E,F) The same as in (A,B) except that *g*_*gap*_ is replaced by the GABA input rate (*g*_*GABA*_). Network activity is generated by setting *g*_*GABA*_ to 0 (E), and 5 (F) nS in response to a fourteen-fold transient increase in granule cell AMPA input at time 0 ms for 2 ms. (G,H) The same as in (C,D) except that *g*_*gap*_ is replaced by *g*_*GABA*_. (I,J) Heatmaps showing the effect of altering both *g*_*gap*_ and *g*_*GABA*_ on mean peak coactivity (I) and mean (ΔPLV) (J). Heatmaps are colour-coded according to colour-bars on top of each panel.

Given the presence of MLI-MLI inhibition in the network, the combined role of this inhibition with AMPA and NMDA inputs is then assessed by altering the conductance of GABAergic current (*g*_*GABA*_). Removal of MLI-MLI inhibition by setting *g*_*GABA*_ = 0 nS causes a loss of global network oscillatory coactivity (Figure 8E), whereas increasing GABAergic inhibition by setting *g*_*GABA*_ = 5 nS induces synchronized activity and lowers elevated coactivity post AMPA input (Figure 8F). With increased inhibition (i.e. higher *g*_*GABA*_), peak coactivity decreases (Figure 8G; Spearman *ρ* = −0.911, p ≤ 0.001) in a manner similar to exclusively applying AMPA input (Figure 6H). However, network synchronization, as measured by ΔPLV, initially decreases when GABAergic inhibition is added, but subsequently increases when *g*_*GABA*_ is increased further (Figure 8H; Spearman *ρ* = 0.840, p ≤ 0.001), suggesting that modest inhibition decreases network synchronization and more prevalent MLI-MLI inhibition promotes the transient NMDA mediated synchronization of the MLI network. These findings are consistent across magnitudes of gap junctional coupling (*g*_*gap*_) and MLI-MLI inhibition (*g*_*GABA*_), with the mean peak coactivity across networks increasing primarily as a result of decreased *g*_*GABA*_, whereas the mean ΔPLV across networks increasing with increasing *g*_*gap*_ (Figure 8I) as well as exhibiting a biphasic profile with increasing *g*_*GABA*_ (Figure 8J).

One important feature that warrants highlighting is that, in the absence of gap junctional coupling (*g*_*gap*_ = 0 nS), increasing MLI-MLI inhibition in the network by increasing *g*_*GABA*_ does not elevate the mean ΔPLV above its value when both gap junctional coupling and MLI-MLI inhibition are absent (i.e. *g*_*gap*_ = 0 nS and *g*_*GABA*_ = 0 nS; Figure 8J) even at large levels of inhibition when *g*_*GABA*_ = 5 nS. This suggests that MLI-MLI inhibition alone is insufficient to generate enhanced network synchronization, but rather augments (or strengthen) existing synchronization within the network. In contrast, increasing gap junctional coupling in the absence of MLI-MLI inhibition produces progressive increase in ΔPLV (Figure 8J), suggesting that gap junctional coupling is the primary driver of the network synchronization and damped oscillatory coactivity, while GABAergic inhibition is subsequently modulating these dynamics. These findings suggest that network synchronization driven by MLI gap junctions is likely eliciting more synchronous MLI-MLI inhibition which in turn enhances network synchronization in a positive feedback loop.

Collectively, these network simulations provide mechanistic insights into the sensory-evoked responses of MLIs detected *in vivo* (Brown et al., 2025). When binning these *in vivo* MLI responses according to maximal coactivity as done in Brown et al. (2025), an initial large collective response followed by elevated activity that exhibits pronounced peak coactivities indicative of network synchronization is observed (Figure 9A). The relationship between peak coactivity and number of peaks detected in the 50 ms window after initial response is further quantified (see Methods; Figure 9B), showing that there is a rise in the number of peaks at higher coactivity level. Repeating the analysis on the MLI network model, we find that increasing AMPA (transient AMPA rate increasing by two-to twenty-fold relative to basal AMPA input rate *λ*_*P, basal*_ in steps of 3 x *λ*_*P, basal*_) and NMDA (*g*_*NMDA*_ increasing from 0.5 to 2.0 nS in steps of 0.25 nS) inputs qualitatively reproduces the *in vivo* activity observed (Brown et al., 2025) (Figure 9C-D). This network model analysis thus predicts that the large synchronous activity observed with 6-8 ms latency after sensory stimulus in MLIs *in vivo* likely reflects a brief increase in glutamatergic AMPA input into MLIs with the subsequent elevated firing resulting from NMDA activation, while the oscillatory coactivity observed after AMPA evoked synchronization is likely a product of gap junction-dependent synchronization and MLI-MLI inhibition in response to the NMDA current.

**Figure 9:**
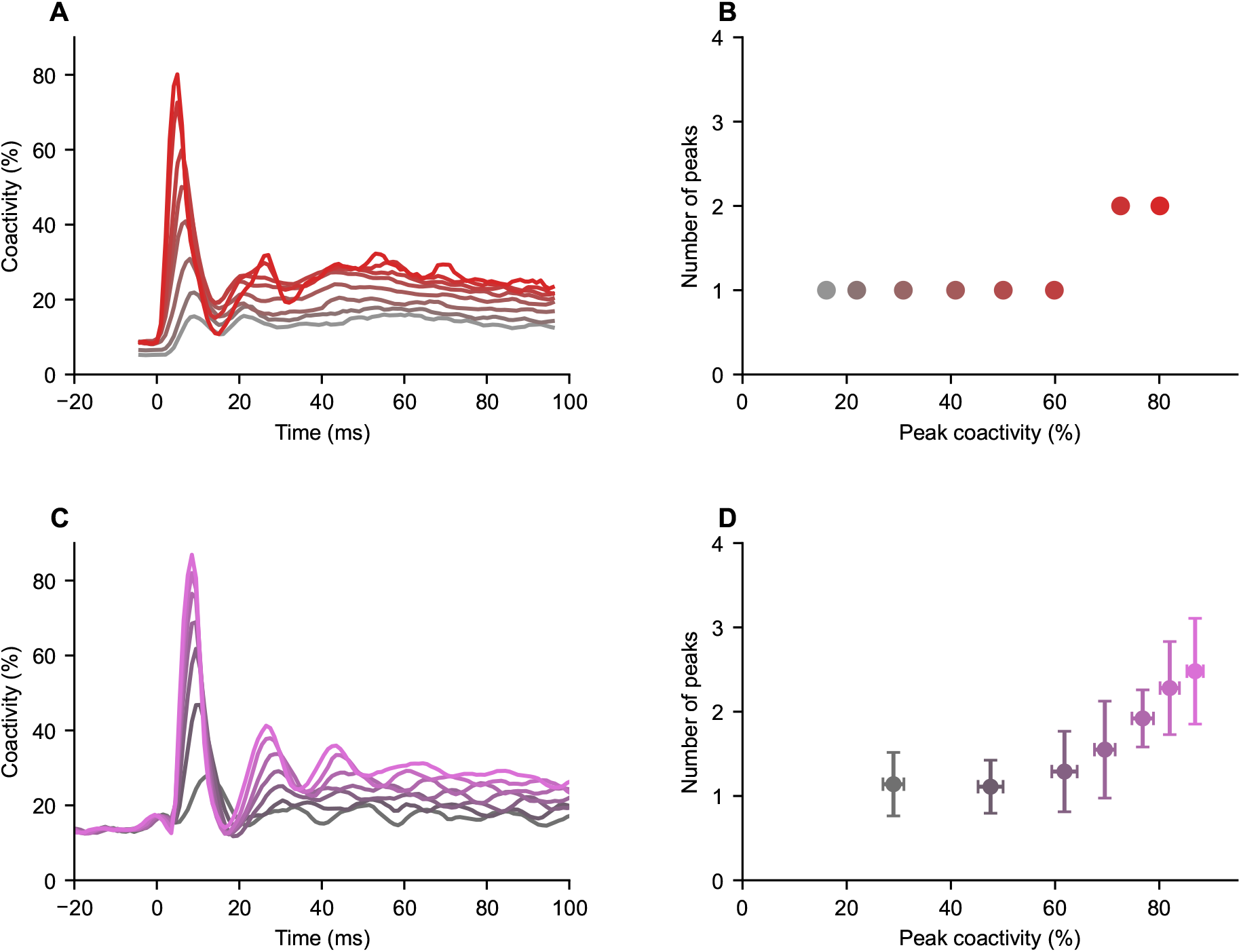
Comparison of MLI coactivity *in vivo* to that generated by MLI network models. (A) Coactivity traces for MLIs recorded *in vivo* in response to a sensory stimulation using voltage imaging by Brown et al. (2025) binned by maximal coactivity. (B) Number of peaks in response to a sensory stimulation as a function of peak coactivity for MLIs recorded by Brown et al. (2025). (C) Coactivity traces generated by a network model MLIs resulting from increasing basal AMPA input rate (*λ*_*P, basal*_) two-to twenty-fold (in steps of 3 x *λ*_*P, basal*_) applied at time 0 ms for 2 ms, and increasing NMDA input (*g*_*NMDA*_ from 0.5 to 2.0 nS in steps of 0.25 nS). (D) Number of peaks in elicited in responses to AMPA and NMDA inputs described in (C) as a function of peak coactivity generated by *n* = 100 realizations of the network model. Model results are shown as mean and standard deviation.

## Discussion

This study uses both cable theory and conductance-based modelling to explore the dendritic filtering and gap junctional coupling of MLIs. Using this framework, we show that predominantly passive dendritic filtering as well as the strength and position of gap junctional coupling dictate gap junction-dependent synchronization in pairs of ball and stick MLI models. Building on these findings, we have further developed a network of gap junctionally coupled one compartment Hodgkin-Huxley type MLI models to investigate the role of gap junctions and MLI-MLI inhibition. These simulations reveal that gap junctional coupling (GABAergic inhibition) drives (augments) transient network synchronization of MLIs in an NMDA-mediated manner, producing responses consistent with MLI activity recorded *in vivo*.

### Active dendrites and gap junctional coupling

Although voltage-gated dendritic conductances have been shown to contribute to synaptic filtering and firing properties of neurons (Migliore and Shepherd, 2002; Tran-Van-Minh et al., 2015; Gollo et al., 2009) along with modulating the effects of gap junctional coupling on network dynamics (Saraga et al., 2006), the MLI modelling in this study suggests that the passive properties of MLI dendrites dictate the effects of gap junctional coupling in MLIs. This finding is consistent with previous studies showing predominantly passive dendritic filtering of synaptic inputs in MLIs (Biane et al., 2021; Tran-Van-Minh et al., 2015; Abrahamsson et al., 2012), but contrasts with others showing supralinear dendritic Ca^2+^ integration in MLIs (Tran-Van-Minh et al., 2015). Asymmetrical gap junctional coupling, such as that produced by gap junctions connecting different dendritic positions, have not been assessed using ball and stick models here. Dendritic location, input resistance, internal dendritic resistance and directional coupling alter interactions between pairs of cells, but can also mask asymmetry in other facets of coupling (Mendoza and Haas, 2022). Such asymmetries can affect the filtering and transmission of gap junctional currents within dendritic compartments, but are unlikely to recruit active dendritic conductances more strongly than large symmetrical, proximal gap junctions investigated here (which we have shown here to result in large voltage deflections in dendritic compartments). As such, it is likely that the predominant effect of passive filtering and the gap junction-dependent synchronization described here with symmetrical gap junctional coupling also extends to asymmetrical gap junctional coupling in MLIs.

### Transient network synchronization of MLIs

Gap junction-dependent transient network synchronization occurs in a number of neuronal networks. For example, gap junctional coupling underlies high frequency network oscillations in the hippocampus, known as ripples (LeBeau et al., 2003; Traub et al., 1999; Draguhn et al., 1998), which get abolished following the application of anesthetics, such as halothane, that block gap junctions (Burt and Spray, 1989; Ylinen et al., 1995). In the cerebellum, synchronous complex spikes in PCs are dependent on gap junctional coupling of inferior olivary neurons (Blenkinsop and Lang, 2006; Llinás and Sasaki, 1989), which give rise to synchronized climbing fibre input into PCs that is time-locked to movement (Welsh et al., 1995). In networks of electrically coupled MLIs (Kim et al., 2014; Mann-Metzer and Yarom, 1999; Alcami and Marty, 2013) like those modelled here, a time resolved transient increase in AMPA input likely homogenizes the action potential phase, increasing the effective *g*_*gap*_ and strengthening the effects of gap junctional coupling transiently. In other words, the combined electrical coupling increases the effective *g*_*gap*_, shifting the dynamics of coupled MLIs into a regime of faster synchronization with substantially lower steady state phase difference (ΔΦ_*ss*_) and synchronization time constant (Figure 4A).

The network modelling performed here reproduces the *in vivo* MLI responses to sensory stimuli (Brown et al., 2025) (Figure 9). However, the utility of the network MLI model is the ability to investigate manipulations that would be difficult to perform *in vivo*. Indeed, the investigation of gap junctional coupling, GABAergic MLI-MLI inhibition strength, and their interplay in the network model uncovers complex network dynamics where slow inputs, such as NMDA activation after glutamate spillover, results in gap junctional mediated increases in network synchrony with MLI-MLI inhibition enhancing unexpectedly synchronous activity within the MLI network. This suggest that the properties of gap junctions as well as network connectivity contribute to MLI activity in the molecular layer *in vivo*.

Functionally, CSCs low-pass filter and BCs high-pass filter parallel fibre inputs to set the frequency band of transmission to PC and to regulate PC gain and firing pattern (Rizza et al., 2021; Masoli et al., 2025). Given that electrical synapses increase the number of MLIs that converge onto a PC (Kim et al., 2014), one would expect NMDA excitation of MLIs to inhibit PC firing *in vivo* (Liu et al., 2014). With the transient gap junction-dependent synchronization demonstrated here during NMDA input, it is likely that the regulation of PC activity by CSCs and BCs is enhanced by electrical synapse synchronization. Notably, NMDA dependent stimulus evoked transient bursting (and spike-adding) has been shown to be tightly regulated by dendritic Ca^2+^ influx and Ca^2+^ activated K^+^ currents in two-compartment conductance-based models of CSCs (Koch et al., 2026). Such mechanism could be also in action in modulating gap junction-dependent transient network synchronization evoked by NMDA input.

Gap junctional coupling in MLIs is likely important to network responses not only during sensory evoked MLI synchrony that drives movement (Brown et al., 2025), but also directly in motor activity. For example, synchronization of pairs of MLIs during saccades in marmosets (Shadmehr et al., 2026) suggests a broader general role of gap junctional coupling in MLI network activity. Additionally, MLI inhibition of PCs has been implicated in cerebellar learning (Wulff et al., 2009; Ma et al., 2020; Bonnan et al., 2022). Gap junctional coupling driven transient synchronization of MLIs, and modulation thereof, may therefore be an important process in cerebellar learning.

### Limitations

The complex dendritic arbor of MLIs are approximated here as a single dendritic cable to enable thorough investigation of the effects of gap junctional coupling on synchronization in pairs of MLIs. Dendritic branching can affect dendritic filtering (Ferrante et al., 2013) and input segregation on multiple dendritic branches may further contribute to dendritic filtering in MLIs, an aspect that is neglected here. However, dominance of passive properties in filtering in the thin MLI dendrites has been previously observed (Biane et al., 2021; Tran-Van-Minh et al., 2015; Abrahamsson et al., 2012) and likely dominate dendritic filtering irrespective of the inclusion of multiple dendritic branches.

Additionally, the full connectivity of MLIs in the cerebellar cortical circuit are not considered in the network simulations performed here. Only parallel fibre excitation of MLI models is considered, although climbing fibres also innervate MLIs (Kim and Augustine, 2021; Szapiro and Barbour, 2007). High frequency parallel fibre (Nahir and Jahr, 2013; Carter and Regehr, 2000b; Clark and Cull-Candy, 2002), concurrent parallel and climbing fibre activity (Szapiro and Barbour, 2007; Jörntell and Ekerot, 2002, 2003) and activation of multiple climbing fibres (Kim and Augustine, 2021) are all associated with glutamate spillover and extrasynaptic NMDA receptor activation. Since the AMPA and NMDA inputs used here are implemented in a source agnostic manner, the effects reported here are likely not parallel fibre specific instead reflecting the distinct timescales of AMPA and NMDA input currents. Thus, the NMDA dependent effects reported here, including the gap junction-driven transient synchronization, are not dependent on the specific source of the glutamate activating NMDA receptors, but rather that sufficient glutamate is released during the sensory evoked response causing NMDA receptor activation. Moreover, the effect of Purkinje cell inhibition of MLIs (Blot et al., 2016; Kim and Augustine, 2021) is also not considered; this additional time delayed inhibition may contribute to the transient synchronization characterized here.

The approximation of dendro-dendritic gap junctional coupling in the modelled MLI network as a low pass filter with a decay determined by the length constant does not account for the possibility of dendritic synchronization that is independent of somatic synchronization due to electrical compartmentalization of distal dendritic compartments. However, the effects of such compartmentalization-driven synchrony are not observed in pairs of gap junctionally coupled ball and stick MLI models across different manipulations of dendritic current conductances.

## Conclusion

Overall, our study highlights the passive dendritic properties of MLIs, the role of gap junctions and chemical synapses in shaping transient sensory stimulus evoked synchronization seen in MLIs *in vivo*. Although gap junctional coupling in MLIs and the synchronizing effects of such electrical coupling has been previously established in MLIs (Kozareva et al., 2021; Sotelo and Llinás, 1972; Mann-Metzer and Yarom, 1999; Hoehne et al., 2020; Kozareva et al., 2021; Han et al., 2018; Kim and Augustine, 2021; Kim et al., 2014; Alcami and Marty, 2013), this is, to our knowledge, the first examination of the involvement of gap junctional coupling in MLI activity *in vivo*. Our results provide a comprehensive analysis of the role of dendritic properties in electrical coupling of MLIs and uncovers its role in transient network synchronization, suggesting a critical role of electrical coupling in the function of MLI networks within the cerebellar cortex.

## Additional Information

### Competing interests

The authors declare no competing interests.

### Author contributions

N.A.K: Conceptualization, Methodology, Software, Validation, Formal analysis, Investigation, Writing – Original Draft, Writing – Review & Editing, Visualization. A. K.: Conceptualization, Resources, Writing – Review & Editing, Supervision, Funding acquisition.

### Funding

This work was supported by the Natural Sciences and Engineering Research Council of Canada (NSERC) Discovery (RGPIN-2019-04520) and Alliance International (ALLRP 588367-23) grants to AK. NAK was supported by the Canada Graduate Research Scholarship – Doctoral program. This research was enabled in part by support provided by Calcul Québec (https://www.calculquebec.ca), BC DRI Group, and the Digital Research Alliance of Canada (https://alliancecan.ca/).

## Notes

### Competing Interest Statement

The authors have declared no competing interest.

