## Supplementary Material for "Gap junctional coupling of molecular layer interneurons enables transient NMDA driven synchronization"

### Supplemental Material

#### Tables

**Table S1:** Probability of chemical synaptic connectivity between molecular layer thirds (lower, middle and upper), where lower corresponds to the most proximal layer relative to Purkinje cell layer and upper is the most distal. Data reproduced from Rieubland et al. (2014), with values scaled by the maximum overall distance-dependent connection probability ($0.245$; Figure S1B) listed in parentheses.

| | | Presynaptic | | |
|---|---|---|---|---|
| | | lower | middle | upper |
| | lower | 0.31 (1.27) | 0.25 (1.02) | 0.13 (0.53) |
| Postsynaptic | middle | 0.10 (0.41) | 0.20 (0.82) | 0.27 (1.10) |
| | upper | 0.0 (0.0) | 0.07 (0.29) | 0.16 (0.65) |

**Table S2:** Probability of electrical synapses between molecular layer thirds (lower, middle and upper), where lower corresponds to the most proximal layer relative to the Purkinje cell layer and upper is the most distal. Data reproduced from Rieubland et al. (2014), with values scaled by maximum overall distance-dependent connection probability ($0.56$; Figure S1B) listed in parentheses.

| | lower | middle | upper |
|---|---|---|---|
| lower | 0.61 (1.09) | — | — |
| middle | 0.46 (0.82) | 0.48 (0.86) | — |
| upper | 0.13 (0.23) | 0.380 (0.68) | 0.5 (0.89) |

### Figures

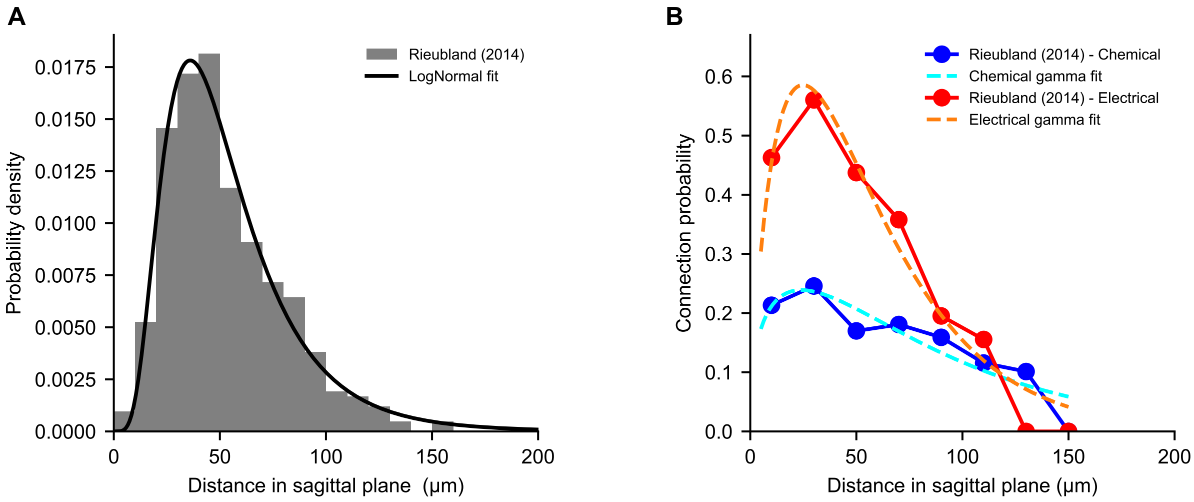


**Figure S1: Intersomatic MLI distances, and chemical and electrical connectivity probabilities**. (A) Lognormal probability density function (see Methods) fit to the distance between MLIs in the sagittal plane quantified by Rieubland et al. (2014). (B) Distance-dependent probability of chemical (blue) and electrical (red) connections reproduced from Rieubland et al. (2014), together with their gamma function fits (see Methods) for chemical (cyan) and electrical (orange) connections.

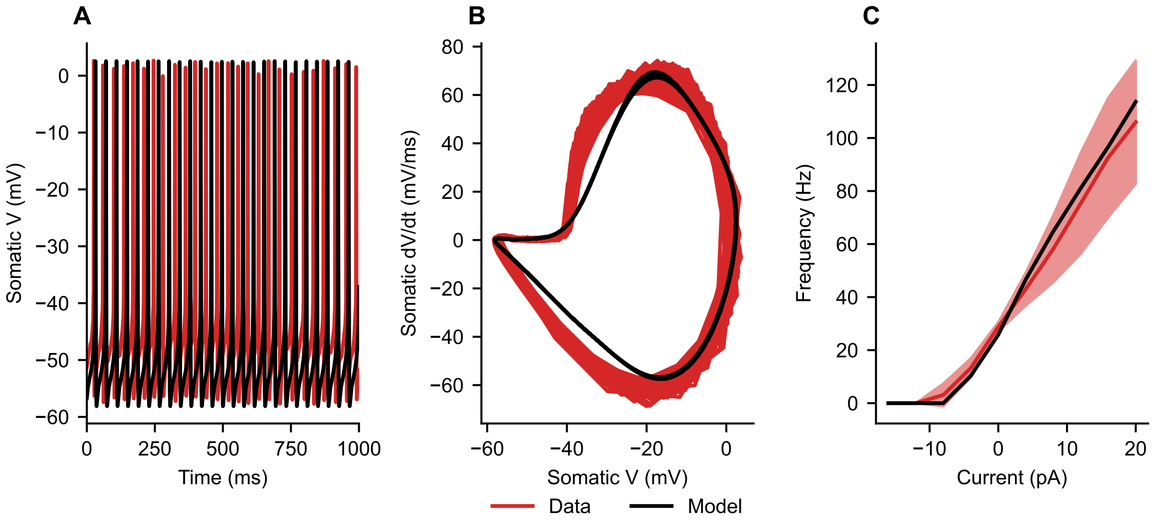


**Figure S2: Fit of the one compartment MLI model**. (A) One compartment model produces spontaneous somatic membrane potential activity (black), without current injection, resembling MLI activity (red) recorded using gap-free patch clamp protocol (Locatelli et al., 2020). (B) Action potential phase cycles in the $(V, dV/dt)$-plane produced by the MLI one compartment model (black) and recorded in CSCs (red). (C) Frequency-current curves generated by the MLI one compartment model (black) and obtained from whole cell patch clamp recordings of CSCs (red) (Locatelli et al., 2020).

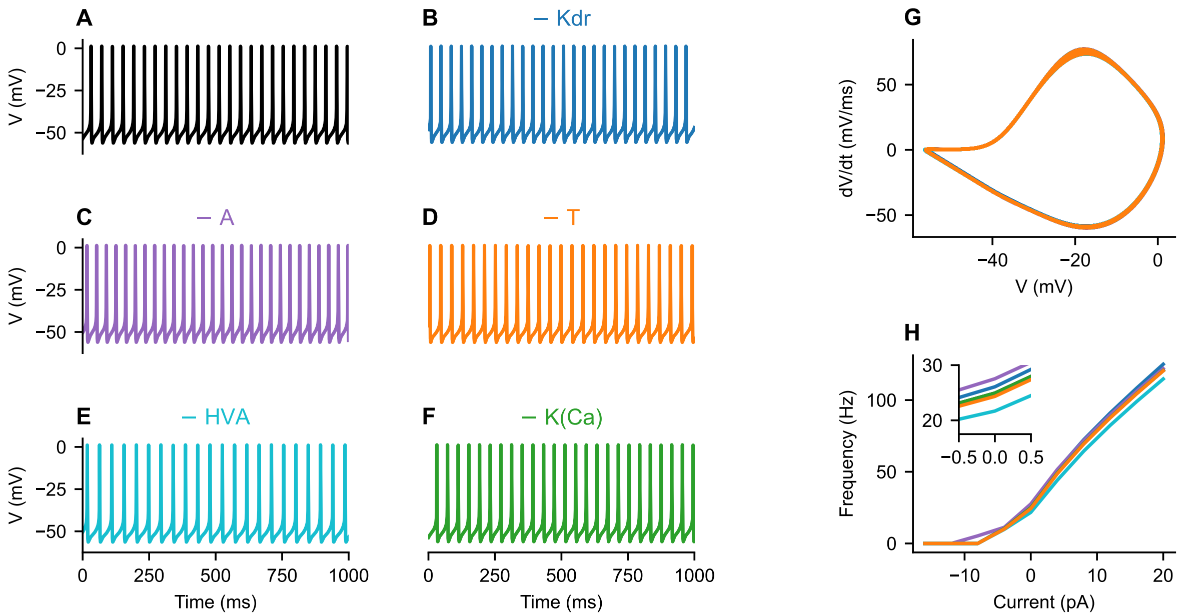


**Figure S3: Dendritic currents in MLI ball and stick model largely do not alter its firing properties**. (A) Firing behaviour of the original MLI model (A), and models in which individual dendritic currents are removed, including delayed rectifier $K^+$ (B; − Kdr), A-type $K^+$ (C; − A), T-type $Ca^{2+}$ (D; − T), high voltage activated $Ca^{2+}$ (E; − HVA) and $Ca^{2+}$ activated $K^+$ (F; − K(Ca)) currents. (G) Action potential phase cycles in the $(V, dV/dt)$-plane produced by the original model as well as models in which individual dendritic currents specified in (A) are removed. (H) Frequency-current curves produced by the original model as well as models in which individual dendritic currents specified in (A) are removed. Inset: The region around zero current input to highlight the small offsets in firing rate with removal of Kdr, A-type, T-type and HVA dendritic currents.

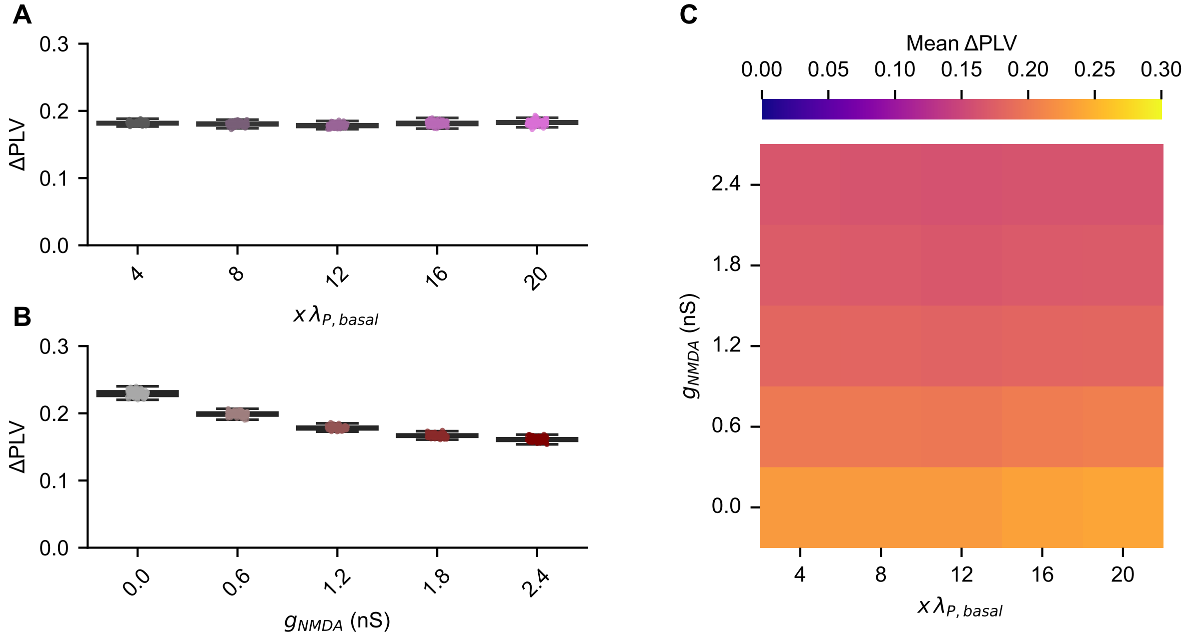


**Figure S4: Removal of gap junctional coupling does not effect AMPA and NMDA-dependent changes of pairwise phase locking in MLI networks**. (A,B) The effect of increasing both basal AMPA input rate ( x $\lambda_{P,\,basal}$) (A; Spearman $\rho = 0.084$, p = 0.06) and NMDA input ($g_{NMDA}$) (B; Spearman $\rho = -0.971$, p $\leq$ 0.001) on ΔPLV in MLI networks lacking gap junctions. (C) Heatmaps showing the effect of altering both $\lambda_{P,\,basal}$ and $g_{NMDA}$ on ΔPLV. Heatmap is colour-coded according to the colour-bar on top of the panel.
